# A Pair of DNA Glucosyltransferases Elevate Counter-defense in Bacteriophage T4

**DOI:** 10.64898/2025.12.06.692756

**Authors:** Luis Ramirez-Chamorro, Frédéric Bonhomme, Anton Lukas Ipsen Wolff, François Lecointe, Marcel Hollenstein, Mart Krupovic, Marianne De Paepe, Yuvaraj Bhoobalan-Chitty

## Abstract

Bacteriophages encode diverse pathways to modify their nucleobases. These modifications help phages to evade the host defense systems such as restriction-modification (RM), and type II and type V CRISPR-Cas systems. On the other hand, modifications can also serve as a target for other host defense systems, illustrating the complexity of the defense and counter-defense landscape. Bacteriophage T4 encodes two glucosyltransferases (GTs), α-GT and β-GT, that post-replicatively add a glucose moiety to the hydroxymethylated deoxycytosines (5-hmC) on phage DNA in the α- and β-conformation, respectively. Among all known phages, only six closely related phages encode both α-GT and β-GT. Here, through biochemical and genetic analysis, we show that β-GT has higher catalytic activity, whereas α-GT is more strongly expressed. During the T4 infection, these factors determine the contributions of both GTs, with α-GT and β-GT contributing respectively to glucosylation of 66% and 33% of all 5-hmC. Encoding a single GTs is sufficient for T4 to overcome the *E. coli* type I and type IV RM systems, unless the glucosylation capacity decreases below the 80% threshold. However, when encountering a host encoding DNA glycosylase Brig1 in addition to type I and type IV RM systems, a second GT is necessary to enable Brig1 escapers to resist RM systems. These results demonstrate that encoding multiple GTs with redundant functionalities provides an evolutionary advantage when simultaneously confronted with multiple antiphage defense systems.

## INTRODUCTION

Bacteriophages, particularly members of the class *Caudoviricetes*, encode a variety of pre-replicative and post-replicative pathways to diversify their DNA nucleobase content beyond canonical composition ^1^. Modifications range from methylation, the most prevalent modification among all domains of life, to complex modifications that result in hypermodified bases such as glucosylmethylcytosine (5-gmC), aminocarboxymethyladenine and queuosine-like 7-deazaguanine derivatives ^2,3^. Methylation is a key component of the host-encoded RM systems that target viruses ^4^. Similarly, hypermodification has been shown to be essential in phage T4 to evade host RM systems, as well as type II and type V CRISPR-Cas systems ^5–10^. Additionally, modifications enable recognition of non-phage (host) DNA by phage proteins, thereby contributing to transcriptional repression ^11^ and cleavage of host DNA by phage endonucleases ^12,13^.

Hypermodified DNA bases, in particular 5-gmC, were first identified among T-even phages ^14^. Phage-encoded enzymes first modify the host nucleotide precursor pool, converting deoxycytidine diphosphate (dCDP) and deoxycytidine monophosphate (dCMP) into hydroxymethylated deoxycytidine triphosphate nucleotides (5-hmdC) ^15^. During phage replication, the modified nucleotides are incorporated into the nascent DNA. Subsequently, the glucosyl group from uridine diphosphate glucose (UDP-glucose) is transferred onto the hydroxymethylated cytosine (5-hmC) by glucosyltransferase enzymes (GTs) ^16^. In phage T4, all cytosines in DNA are exclusively mono-glucosylated by primary glucosyltransferases, either alpha-glucosyltransferase (α-GT) or beta-glucosyltransferase (β-GT) ^17,18^. Among these, 60% possess α-glycosidic linkages (catalysed by α-GT) while 40% have β-glycosidic linkages (catalysed by β-GT) ^19^. In T4, it has been suggested that glucosylation by α-GT is concomitant with replication and occurs upon association of the enzyme with the T4 replisome, whereas the remaining non-glucosylated 5-hmdC residues serve as substrates for β-GT ^20^. In phages T2 and T6, that possess only the primary glucosyltransferase α-GT, β-glucosyl-HMC-α-GT acts as a secondary glucosyltransferase, catalysing the addition of a glucosyl group onto an already α-glucosylated cytosine, resulting in 5-gentiobiosyl-methylcytosine base ^17,18,21,22^.

The diversification of modifications likely results from an evolutionary arms race between phages and their hosts. In the *E. coli*–T4 host-phage system, hydroxymethylation of cytosine protects the phage DNA from the widely conserved type I EcoKI RM system ^23^. Subsequent glucosylation protects from the methylation specific type IV McrBC/McrA RM systems ^5,24^. DNA glycosylases such as Brig1 and the prophage encoded type IV RM system GmrSD (glucose-modified restriction) are both capable of antiphage activity by targeting 5-gmC ^25–27^. Brig1 acts specifically against α-glucosylated cytosine, whereas GmrSD targets both stereoisomers of 5-gmC and the 5-gentiobiosyl-methylcytosine. The activity of GmrSD and its variants is inhibited by phage T4 proteins IPI, IPII and IPIII ^25,26,28^. As an alternative to glucosylation, arabinosylation of the hydroxy deoxycytosine (5-hdC) in the phage DNA ^29,30^ also offers robust protection not only against type II RM and the DNA glycosylase Brig1 ^31^, but also against type I CRISPR-Cas. In addition to Brig1, antiviral glycosylases such as Dag1/Dag2 and Brig2, which cleave modified guanine and thymine bases, respectively, have also been characterized ^32^.

Our study reveals that while most phages encode only one primary GT, a few closely related phages, including phage T4, encode both α-GT and β-GT. The two phage GTs appear to have diverged from a common cellular ancestor with the GT-B structural fold following a gene duplication event, resulting in paralogous enzymes that yield stereochemically different reaction products. We show that the balanced levels of α-and β-glucosylation observed in wild-type phage is due to the opposing strengths of the two GTs, reflecting differences in both their enzymatic activity and gene expression. We demonstrate that each GT, when expressed alone, is sufficient to ensure more than 80% cytosine glucosylation, a level sufficient to provide complete protection against *E. coli* type IV restriction-modification systems. However, despite the apparent functional redundancy of the two GTs in protection against typical RM systems, we show that in the presence of the additional glycosylase-based defense system Brig1, the non-targeted GT serves as an efficient back-up allowing the survival of phage escaper mutants with inactivated target GT.

## RESULTS

### Co-occurrence of two primary glucosyltransferases is rare in phage genomes

Genetic redundancy is rare in phage genomes, due to a high constraint on maximal genome content ^33–35^. Yet β-GT in phage T4 was previously described to have a redundant function with ⍺-GT, which is intriguing ^36^. To investigate this phenomenon, we started by analysing the distribution of GTs (primary and secondary) in phage genomes. We first identified all viral homologs of T-even phage GTs in the RefSeq database by PSI-BLAST (<u>Table S1</u>). The hits were then associated to their respective phage genomes (Figure 1A and <u>Table S1</u>). Among the 111 phages encoding GT homologs, 56 phages carried two types of GT. In all cases, except phage T4 and Bacillus phage G, phage genomes contained a primary-GT alone or in association with a homolog of a secondary-GT. Bacillus phage G contains a second copy of ⍺-GT, whereas phage T4 is unique in encoding two primary GTs, ⍺-GT and β-GT.

**Figure 1:**
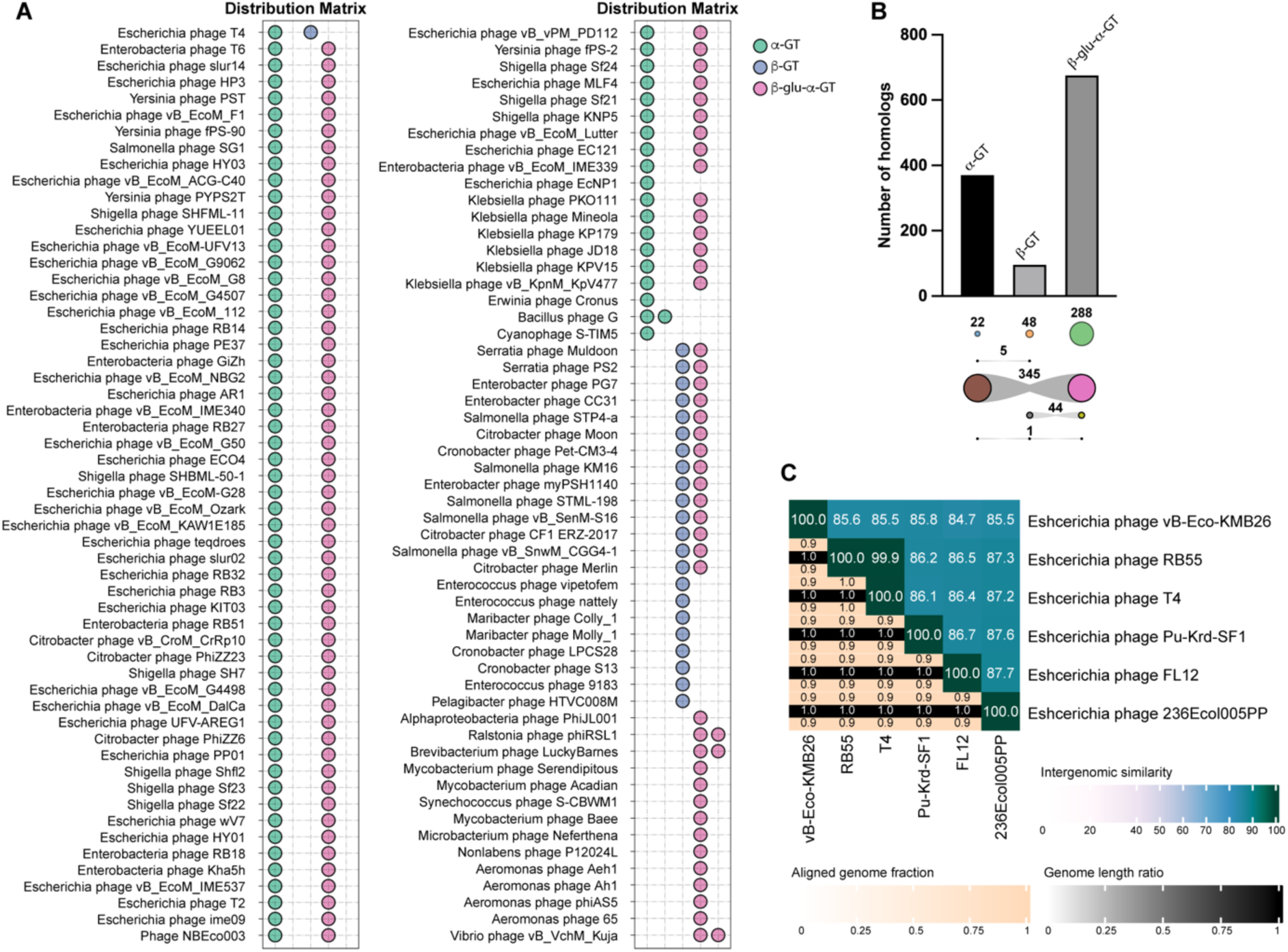
Distribution of glucosyltransferases among phage genomes. **A**. Distribution of α-GT, β-GT and β-glucosyl-HMC-α-GT homologs in phage genomes from the RefSeq database. **B**. Bar-plot showing the total number of α-GT, β-GT and β-glucosyl-HMC-α-GT homologs in viruses (taxid:10239), in the non-redundant (nr) NCBI GenBank database. The number of GT homologs encoded either individually or in combination with other GT(s) is indicated below the x-axis. **C**. Heatmap showing pairwise sequence similarities between the six Escherichia phages that encode both α-GT and β-GT. The top-right half of the heatmap, above the diagonal, displays the percentage of similarity between each genome pair. In the bottom-left half, three values are provided for each pair: the top and bottom values represent the aligned fraction for the phage genome pair corresponding to that row and column respectively. The middle value is the ratio of their genome lengths.

To check if the co-occurrence of two primary-GTs in T4 is an exception, we expanded our analysis to all phage genomes in the non-redundant (nr) NCBI GenBank database. Among the 753 phages that carried at least one homolog of T-even phage GTs, only one encoded all three GT types, and five phages, including phage T4, encoded both primary GTs (Figure 1B and <u>Table S1</u>). The overall nucleotide identity between the six phage genomes ranged from 84% to 88%, with a coverage of 90-100% (Figure 1C), indicating that these phages are closely related and belong to the same *Tequatrovirus* genus. A whole-genome tree showed that the five phages encoding both primary GTs share a common ancestor (Figure S1). This analysis also confirmed that the vast majority of phage genomes carry a single primary GT together with a secondary GT. Surprisingly, several phages also encoded distantly related secondary GTs in the absence of any recognizable primary GT (Figure S2). Collectively, these findings indicate that the co-occurrence of ⍺-GT and β-GT is an uncommon event which may indicate either that it does not confer a selective advantage to the phage or that the acquisition of the second primary GT is a relatively recent event in phage evolution.

### Evolutionary relationship between α-GT, β-GT and β-glucosyl-HMC-α/β-GT

To investigate this hypothesis, we determined the evolutionary relationships between the three GT types. Homologs of the ⍺-GT, β-GT and β-glucosyl-HMC-α/β-GT do not display any appreciable sequence similarity across the GT types (<u>Table S1</u>). However, structural comparison showed that ⍺-GT and β-GT adopt the same structural fold ^37,38^, known as the GT-B ^39^ (Figure 2), suggesting that the two enzymes are evolutionarily related. By contrast, analysis of the predicted structural model of β-glucosyl-HMC-α/β-GT from phage T6 generated with AlphaFold3 revealed that it belongs to an evolutionarily distinct superfamily of GTs that adopt the GT-A fold ^39^ (Figure 2). To gain further insights into their origins and evolution, we performed structural clustering of the phage GTs with representatives from diverse GT families using DALI ^40^. ⍺-GTs and β-GTs represent two distinct GT families, GT63 and GT72, respectively, which are exclusively found in phages ^41^. However, our analysis showed that in structural comparisons, the two families clustered together, forming sister groups in the structure-based dendrogram (Figure 2). Notably, among other GT-B superfamily members, the phage GTs were most closely related to the bacterial GT113-family GTs (Figure 2). These results suggest that ⍺-GT and β-GT have evolved following gene duplication from a common ancestor, which itself has originated from a cellular GT. β-glucosyl-HMC-α/β-GT homologs from phages clustered together as a sister group to the GT2 family GTs of the GT-A fold superfamily, also suggesting cellular ancestry of the phage β-glucosyl-HMC-α/β-GTs (Figure 2).

**Figure 2:**
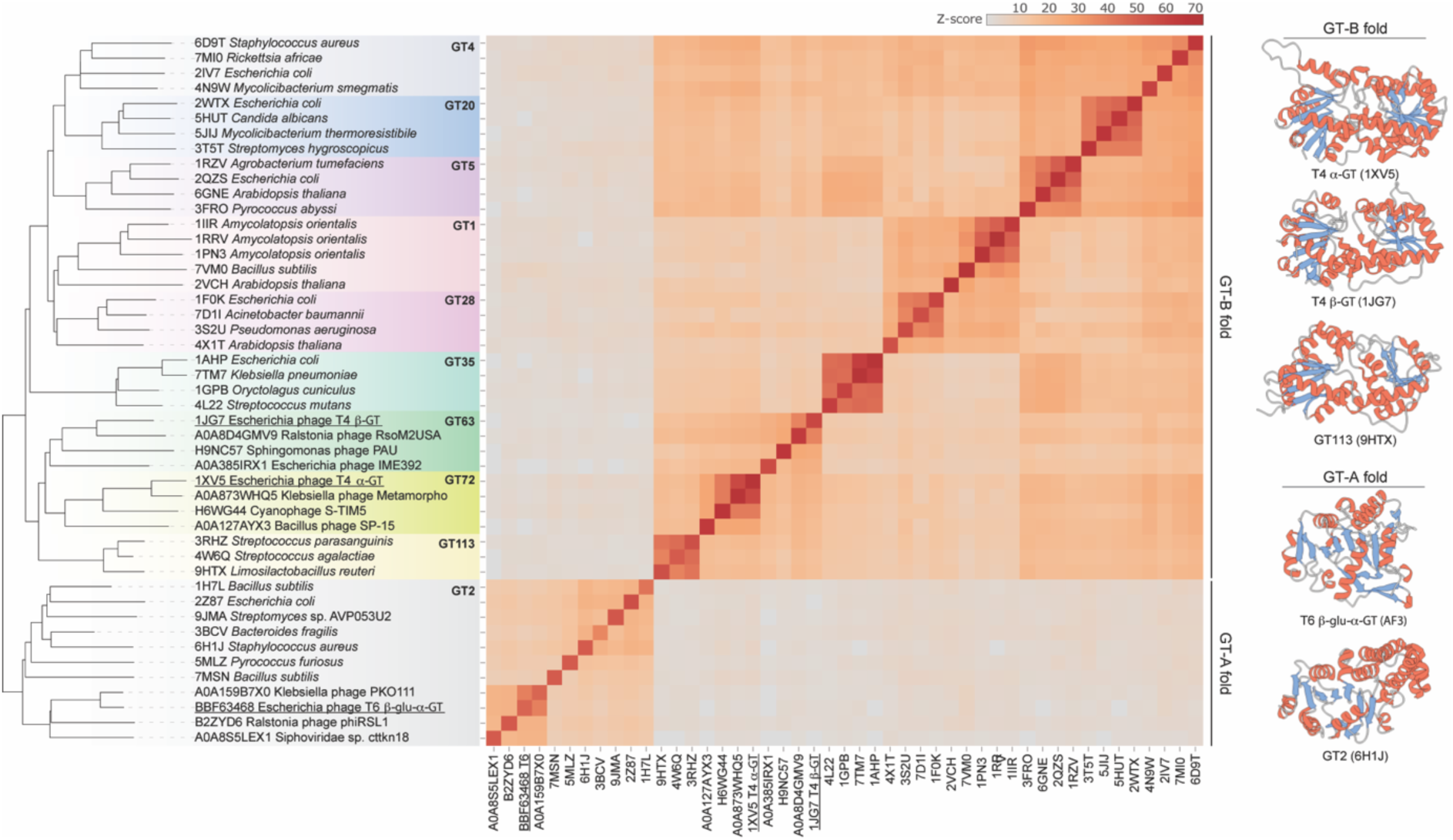
Relationships between α-GT, β-GT and β-glucosyl-HMC-α/β-GT. A dendrogram and heatmap based on pairwise Z-scores, obtained using DALI, comparing T4 α-GT, T4 β-GT and modelled structure of β-glucosyl-HMC-α/β-GT along with representative cellular GTs from diverse families of the GT-A and GT-B superfamilies. Experimentally determined structures and structural models are indicated with the corresponding PDB and UniProt accession numbers, respectively. α-GT and β-GT of phage T4 and β-glucosyl-HMC-α/β-GT of phage T6 are underlined. Different GT families are highlighted with different background colors on the dendrogram. The color scale indicates the corresponding Z scores. On the right, the crystal structures of T4 α-GT (PDB: 1XV5), T4 β-GT (PDB: 1JG7) and the modelled structure of β-glucosyl-HMC-α/β-GT (AF3) are compared to the structures of closest cellular representatives from the GT-B and GT-A fold superfamilies, GT113 (PDB: 9HTX) and GT2 (PDB: 6HI1J), respectively. Structural models are colored according to the secondary structure elements: α-helices, red; β-strands, blue; random coil, grey.

Together, these results suggest that cellular GTs have been coopted for hypermodification of the phage DNA at least twice independently (⍺-GT/β-GT and β-glucosyl-HMC-α/β-GTs, respectively). In the case of the primary ⍺-GT and β-GT, it is tempting to speculate that specificity towards the DNA substrate has evolved once, in the common ancestor of the two families, followed by gene duplication and diversification of the paralogs towards different stereospecificities.

### Analysis of protein-protein interactions associated with α-GT and β-GT

Before addressing the potential advantage of encoding two GTs with different stereospecificities, we investigated the mechanisms behind the distribution of ⍺- and β-glycosidic modifications on cytosines in T4 DNA. Previously it has been estimated that ⍺-GT and β-GT contribute to 60% and 40% of glucosylation, respectively^17,19^. To explain that distribution, it has been previously hypothesized, based on *in vitro* results ^20^, that T4 ⍺-GT binds to gp45, the sliding clamp protein that forms the basis of the T4 replication machinery, and modifies the T4 DNA concomitantly with replication, with β-GT presumed to intervene latter, on unmodified bases. To investigate this possibility, we analyzed the interaction between the sliding clamp protein and ⍺-GT or β-GT. C-terminally histidine tagged versions of ⍺-GT and β-GT were individually purified from *E. coli*. Based on the elution volume during size-exclusion chromatography it was estimated that both proteins exist as monomers (Figure S3). Tag-free gp45 was overexpressed in *E. coli* (Figure S4). Soluble fractions containing either tagged ⍺-GT or β-GT were combined with the fraction containing gp45 prior to immobilized metal affinity chromatography (IMAC) purification. Compared to the controls from which the gp45-containing fraction was omitted, no co-purification of protein matching the size of gp45 was observed in the eluted fractions (Figure 3A), contrary to what was previously observed ^20^.

**Figure 3:**
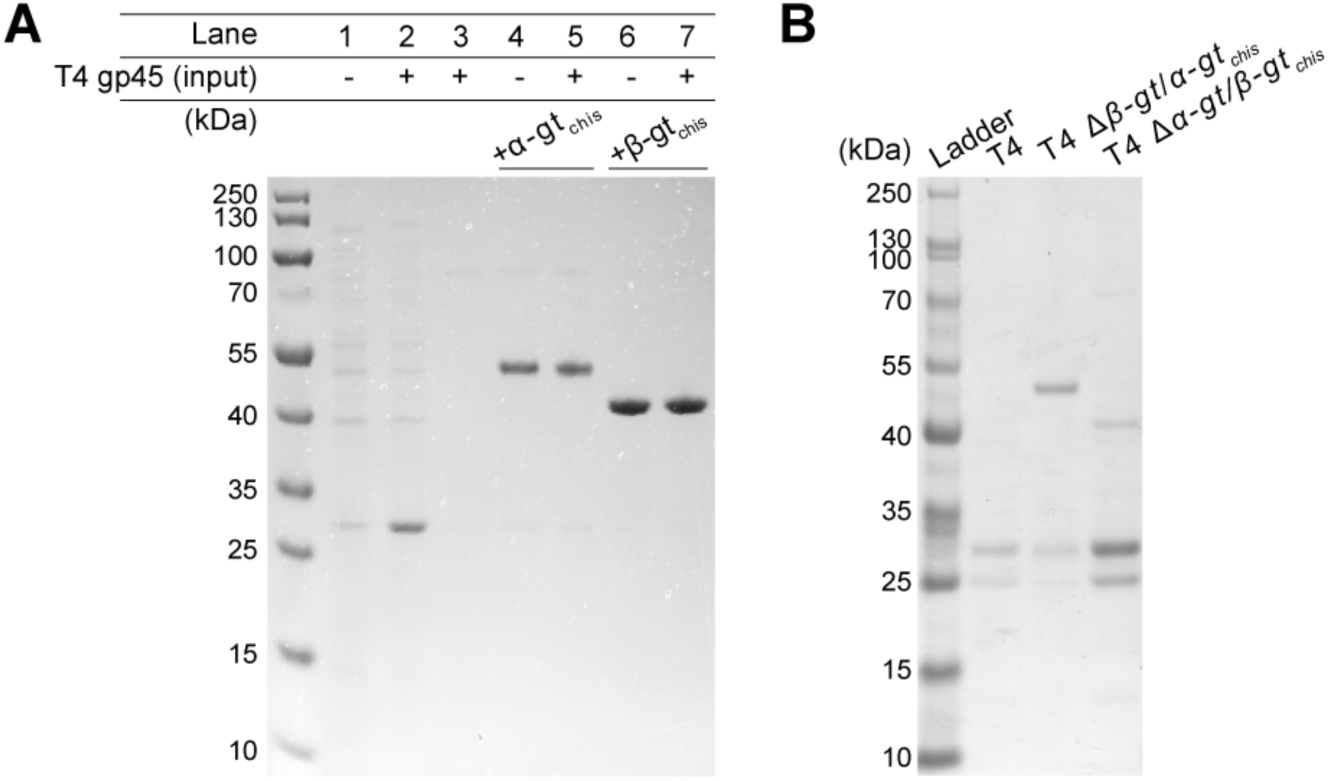
Protein-protein interaction analysis of α-GT and β-GT. **A**. *In vitro* pull-down of histidine tagged α-GT or β-GT, mixed with un-tagged T4gp45. Lane 1 and lane 2 correspond to the total cell extract of *E. coli* BL21(DE3) carrying plasmid-borne un-tagged T4 gp45, before and after induction with IPTG. Lane 3 is a control IMAC purification of un-tagged gp45. Lanes 4 and 5 are IMAC purification of C-terminal histidine tagged α-GT without or with fraction containing tag-free gp45 (shown in lane 2). Lanes 6 and 7 are IMAC purification of C-terminal histidine tagged β-GT without or with fraction containing tag-free gp45 (show in lane 2). **B**. *In vivo* pull-down of α-GT(chis) (lane 2) and β-GT(chis) (lane 3), based on phage-encoded endogenous expression, after infection of *E. coli* MG1655 with the respective phage T4 mutants. Lane 1 is IMAC purification control from *E. coli* MG1655 infected with the wild-type phage.

To identify any possible interactions *in vivo*, we engineered two phages encoding either ⍺-GT or β-GT, along with an in-frame coding sequence for the histidine-tag at their C-termini, T4 Δ*β-gt/⍺-gt_Chis_* and T4 Δ*⍺-gt/β-gt_Chis_* (Figure S5). The recombinant phages did not display any loss of virulence in *E. coli* MG1655, which encodes the type IV RM systems (Figure S6). The protein pull-down experiments were performed at least twice using IMAC purification and analyzed by mass spectrometry-based proteomics. Under *in vivo* conditions, neither ⍺-GT nor β-GT co-purified with the sliding clamp protein or any other phage protein (Figure 3B). Proteomic analysis of each pull-down sample further confirmed the lack of interaction between the two GTs and other phage proteins (<u>Table S2</u>).

Here, through *in vitro* and *in vivo* experiments, we show that neither ⍺-GT nor β-GT interact with the sliding clamp protein or other components of the replication machinery of phage T4, contrary to the published results. This suggests that ⍺-GT and β-GT compete directly for their substrate on T4 DNA.

### Relative enzymatic activity and expression strength govern the distribution of glycosidic linkages

In the absence of interaction with the replisome, this distribution could result from competition between the two enzymes, with ⍺-GT being either produced more, and/or enzymatically more active.

In order to identify the contributing factors underlying the specific glucosylation levels, we estimated the difference in enzymatic activity between the two GTs. Using the C-terminally histidine tagged GTs purified earlier, we performed a competition assay. The two GTs were mixed at different molar ratios and incubated with a 5-hmC containing PCR fragment. The contribution of each GT was estimated from the percentage 5-hmC residues that were either ⍺- or β-glucosylated. In the absence of β-GT, ⍺-GT contributed to the glucosylation of 64% of 5-hmC bases. Upon addition of β-GT, even at a concentration four times lower than ⍺-GT, 91% of 5-hmC were β-glucosylated, with the ⍺-GT contributing to only 0.65% of glucosylations (Figure 4A). These results demonstrate that under the *in vitro* conditions used, β-GT exhibits much higher enzymatic activity than ⍺-GT, despite contributing to only 40% of all glucosylations *in vivo*. This strongly suggests that the higher level of glucosylation by ⍺-GT in T4 phage DNA is not due to its higher enzymatic activity.

**Figure 4:**
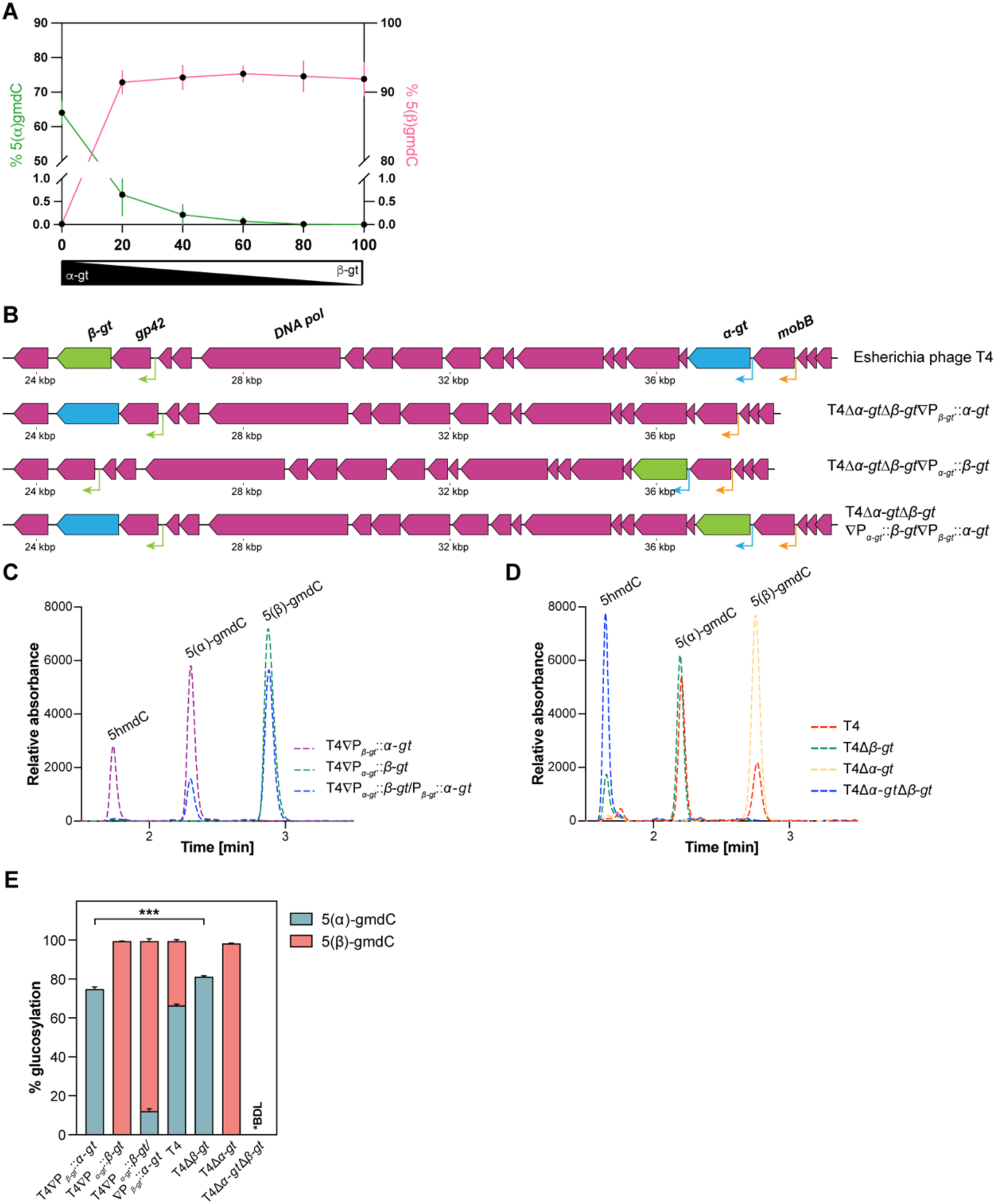
Molecular mechanism behind the competitive glucosylation by α-GT and β-GT. **A**. Competition assay to estimate the relative rate of glucosylation by α-GT and β-GT on a fragment of DNA containing 5-hmdC. The average percentage of 5(α)-gmdC and 5(β)-gmdC nucleotides were estimated by mass-spectrometry. The values shown are average of 6 independent experiments, represented as mean ± SD. **B**. Illustration of phage mutants with genomic location of *α-gt* and *β-gt* swapped between their native locations in the wild-type T4 phage. **C.** LC-MS/MS profile of the mutant phages, T4 ΔP*_α-gt_*::*β-gt*, T4 ΔP*_β-gt_*::*α-gt* and T4 ΔP*_α-gt_*::*β-gt*/P*_β-gt_*::*α-gt* propagated in *E. coli* DH10B. The curves show the abundance of 5-hmdC, 5(α)-gmdC and 5(β)-gmdC nucleotides. **D.** Mass-spectrometric LC-MS/MS profile of the mutant phages, T41′*α-gt*, T41′*β-gt* and T41′*α-gt*1′*β-gt*, in comparison to the wild-type T4 phage. All phages were propagated in *E. coli* DH10B prior to estimation of their nucleotide composition. The curves show the abundance of 5-hmdC, 5(α)-gmdC and 5(β)-gmdC nucleotides. **E**. Average percentage of 5(α)-gmdC and 5(β)-gmdC in phages T4 ΔP*_α-gt_*::*β-gt*, T4 ΔP*_β-gt_*::*α-gt*,T4 ΔP*_α-gt_*::*β-gt*/P*_β-gt_*::*α-gt*,T4, T4 1′*α-gt*, *T4* 1′*β-gt* and T4 1′*α-gt*1′*β-gt*, as estimated by mass spectrometric analysis. The values shown are from three biological replicates, represented as mean ± SD. A two-tailed unpaired t-test of the percentage values of 5(α)-gmdC was used to calculate the *P* values; \*\*\**P* < 0.001.

We then reasoned that the difference could stem from higher or earlier expression of *⍺-gt* during the T4 lytic cycle. An analysis of previously published transcriptomics data ^42^ showed no significant differences in the expression levels of the two GTs, despite slight differences in the timing of expression (Figure S7). The limited difference observed could not conclusively explain the differences in glucosylation levels observed *in vivo*, especially given the significantly higher enzymatic activity of β-GT observed *in vitro*. To elucidate the role of expression strength, we set out to modify the transcription levels of *⍺-gt* and *β-gt*. Due to the technical challenges associated with direct promoter modification on phage genome and lack of knowledge on the ensuing impact on expression strength, we opted instead to switch the positions of the two GTs. We constructed the mutant phage T4ΔP*_⍺-gt_*::*β-gt*ΔP*_β-gt_*::*⍺-gt*, encoding both GTs but under the control of each other’s regulatory sequences, along with two additional phage mutants T4ΔP*_β-gt_*::*⍺-gt* and T4ΔP*_⍺-gt_*::*β-gt*, encoding either ⍺-GT or β-GT under the control of the promoter of the other GT (Figure 4B, Figure S8 and Figure S9, see Methods section).

We then evaluated the impact of the switching of promoters on the relative levels of ⍺-and β-glucosylated cytosine residues by LC-MS/MS. In phage T4ΔP*_β-gt_*::*⍺-gt*, 75% of 5-hmC were glucosylated, exclusively by ⍺-GT (Figure 4C and Figure 4E). By contrast, in phage T4ΔP*_⍺-gt_*::*β-gt*, nearly all 5-hmC were β-glucosylated (Figure 4C and Figure 4E). In T4ΔP*_⍺-gt_*::*β-gt*ΔP*_β-gt_*::*⍺-gt*, 100% of 5-hmC were glucosylated, but only 12% were ⍺-glucosylated, with the remaining 87% being β-glucosylated (Figure 4C and Figure 4E). To interpret these results, we also determined the percentage of glucosylation in the wild-type T4, in the single *gt* deletion mutants that we previously constructed ^43^ and in a double *gt* deletion mutant that was generated in this study (Figure S10). In wild-type T4, 66% of all 5-hmC bases were α-glucosylated and the remaining 33% were β-glucosylated (Figure 4D and Figure 4E), in close agreement with the previous estimations ^17,19^. In the phage lacking both GTs, as expected, all cytosines were only hydroxymethylated. Glucosylation by α-GT increased from 66% in the wild-type to 80% in T4Δ*β-gt*. Similarly, in T4Δ*α-gt* the percentage of glucosylation by β-GT is increased, reaching 99%, while it is only 33% in the wild-type T4.

The increased β-glucosylation when *β-gt* is expressed from P*_⍺-gt_* compared to P*_β-gt_* promoter (88% in phage T4ΔP*_⍺-gt_*::*β-gt*ΔP*_β-gt_*::*⍺-gt* as compared to 33% in wild-type T4) suggests that the expression strength of P*_⍺-gt_* is greater than that of P*_β-gt_*. Further supporting this conclusion, upon expression from P*_β-gt_*, ⍺-GT was able to glucosylate only 75% of 5-hmdC as compared 80% when expressed from its native promoter in phage T4Δ*β-gt*, a small but significant difference. Moreover, concomitant expression of *β-gt* from the P*_⍺-gt_* promoter in phage T4ΔP*_⍺-gt_*::*β-gt*ΔP*_β-gt_*::*⍺-gt* lowers the contribution of ⍺-GT by 7-fold.

Together, these results demonstrate that the *in vitro* enzymatic activity of β-GT is greater than that of ⍺-GT, confirmed *in vivo* by the complete glucosylation of 5-hmC by β-GT in T4Δ*α-gt* in comparison to only 80% glucosylation by ⍺-GT in T4Δ*β-gt*. In phage T4, the higher enzymatic activity of β-GT is compensated by the stronger upstream regulatory sequence and the Shine-Dalgarno (SD) motif associated with *⍺-gt*, resulting in ⍺-glucosylation being the predominant modification.

### Each individual GT is sufficient to provide wild-type level virulence

To investigate the potential selective advantage of encoding two primary GTs, as a first step, we characterized the virulence of T4 GT deletion mutants in relation to their percentage of cytosine glucosylation. To this end, we estimated the killing activity of the wild-type, single and double deletion mutant phages by determining their virulence index (Vi), a recently proposed quantitative measure of the virulence of a phage against a given host ^44^.

Except for the double deletion mutant, T4Δ*α-gt*Δ*β-gt*, the WT and single deletion mutants were able to inhibit growth of the host at all MOIs (Figure 5A). In congruence, Vi estimated over 7 hours of growth post infection was similar between these phages, but the double mutant had a much lower Vi (Figure 5B and Figure S11).

**Figure 5:**
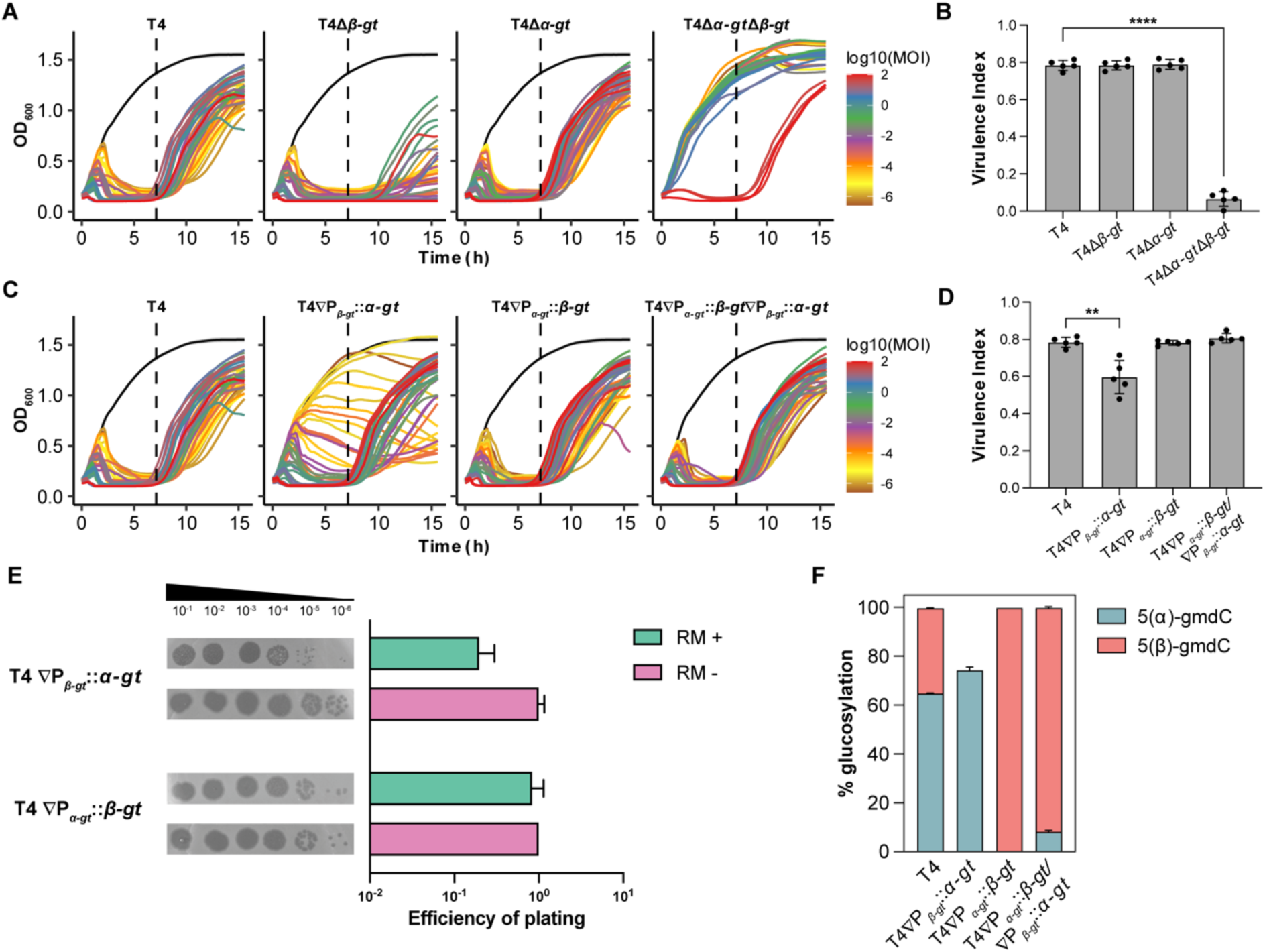
Virulence estimation of phages T4 wild-type, T4 1′*α-gt*, *T4* 1′*β-gt* and T4 1′*α-gt*1′*β-gt* in *E. coli* MG1655. **A**. Growth curves of *E. coli* MG1655 infected with phages T4, T41′*α-gt*, T41′*β-gt* and T41′*α-gt*1′*β-gt*, over a range of MOIs between 10^2^ to 10^−6^. Uninfected culture is represented in black. **B**. Virulence index of the respective phages, as calculated from the growth curves shown in (**A**), up to 7 hours post-infection. Data shown are mean of four biological replicates, represented as mean value ± SD. A two-tailed unpaired t-test of the virulence index values was used to calculate the *P* values; \*\*\*\**P* < 0.0001. **C**. Growth curves of *E. coli* MG1655 infected with phages T4 ΔP*_α-gt_*::*β-gt*, T4 ΔP*_β-gt_*::*α-gt* and T4 ΔP*_α-gt_*::*β-gt*/P*_β-gt_*::*α-gt* and the wild-type T4, over a range of MOIs between 10^2^ to 10^−6^. Uninfected culture is represented in black. **D**. Virulence index of the respective phages, as calculated from the growth curves shown in (**c**), up to 7 hours post-infection. Data shown are mean of four biological replicates, represented as mean value ± SD. A two-tailed unpaired t-test of the virulence index values was used to calculate the *P* values; \*\**P* < 0.01. **E**. Spot assay of serially diluted phages, T4 ΔP*_α-gt_*::*β-gt* and T4 ΔP*_β-gt_*::*α-gt* on *E. coli* DH10B (RM-) or *E. coli* MG1655 (RM+) background. Efficiency of plating (EOP) of phage infection, as estimated from the spot assay, are from four biological replicates, represented as mean ± SD. **F.** Average percentage of 5(α)-gmdC and 5(β)-gmdC in phages T4, T4ΔP*_α-gt_*::*β-gt*, T4ΔP*_β-gt_*::*α-gt* and T4ΔP*_α-gt_*::*β-gt*ΔP*_β-gt_*::*α-gt*, as estimated by mass spectrometric analysis. All phages were propagated in *E. coli* MG1655 prior to estimation of their nucleotide composition. The values shown are from two biological replicates, represented as mean ± SD.

To evaluate the impact of the glucosylation levels obtained with switched promoters on phage virulence, growth of the three mutant phages on *E. coli* MG1655 was compared to that of the wild-type phage (Figure 5C). While the mutant phages T4ΔP*_⍺-gt_*::*β-gt* and T4ΔP*_⍺-gt_*::*β-gt*ΔP*_β-gt_*::*⍺-gt* demonstrated the same levels of virulence as the wild-type phage, the mutant phage T4ΔP*_β-gt_*::*⍺-gt* showed decreased virulence, evident at lower MOIs (Figure 5D and Figure S12). Spot assay of the mutant phages T4ΔP*_⍺-gt_*::*β-gt* and T4ΔP*_β-gt_*::*⍺-gt* on *E. coli* DH10B (no RM systems) and *E. coli* MG1655 confirmed that the decrease in virulence of the latter mutant phage was specific to the presence of RM systems. In addition to the reduced plaque size, a 5-fold decrease in the efficiency of plating was observed with phage T4ΔP*_⍺-gt_*::*β-gt* in the presence of RM systems (Figure 5E).

In light of the glucosylation levels observed in the single *gt* deletion mutants, the decreased virulence observed in phage T4ΔP*_β-gt_*::*⍺-gt* suggests that 80% glucosylation, observed in T4Δ*β-gt*, is the absolute threshold below which phage DNA is susceptible to the activity of the *E. coli* type IV RM systems. Previous estimations of glucosylation level were performed using genomic DNA obtained from phages propagated in an RM-negative strain (*E. coli* DH10B). To determine if the presence of the RM system influences the glucosylation level, the analysis was repeated for phages T4, T4ΔP*_β-gt_*::*⍺-gt*, T4ΔP*_⍺-gt_*::*β-gt* and T4ΔP*_⍺-gt_*::*β-gt*ΔP*_β-gt_*::*⍺-gt*, after propagation in *E. coli* MG1655 (RM-positive) strain. Among the four phages, the percentage of 5-hmC that were α-glucosylated or β-glucosylated remained the same to that observed earlier (Figure 5F and Figure 4E). These results further support that the decreased virulence observed in phage T4ΔP*_β-gt_*::*⍺-gt* was indeed due to a decrease in the percentage of overall glucosylation.

Taken together, these results demonstrate that T4 α-GT and β-GT have evolved to be able to individually glucosylate at least 80% of all 5-hmC bases in the T4 genome, which is sufficient for complete protection from 5-hmC targeting type IV RM systems and is therefore not the explanation for the presence of two GTs. The reduced virulence of phage T4ΔP*_β-gt_*::*⍺-gt*, observed in RM-positive strain, reinforces our previous conclusion that the reduced β-glucosylation in wild-type T4 is due to the greater expression of P*_⍺-gt_* in relation to P*_β-gt_*.

### β-GT is essential for phage T4 survival in wild-type *E. coli* encoding RM and Brig1 defense systems

Despite the redundancy ensuring complete protection against type IV RM systems in the event of loss of a GT, we reasoned that encoding two GTs could be beneficial against other defense systems targeting glycosylated DNA. A DNA glycosylase-based host defense system, Brig1, was recently shown to provide immunity against T-even phages ^27^ by generating abasic sites in DNA containing 5-gmC with ⍺-glycosidic linkages. The authors showed that phage T4 was able to produce escaper mutants in the presence of cosmid-borne Brig1, in the *E. coli* EC100 strain devoid of the type IV RM systems targeting 5-hmC (Figure S13).

To understand the influence of varying levels of ⍺-glucosylation on the anti-phage activity of Brig1, we determined the escaper frequency (⍺-gt inactivating mutants) on a RM-negative strain across a range of phage mutants with varying levels of ⍺-glucosylation. Among phages T4Δ*β-gt* and T4ΔP*_β-gt_*::*⍺-gt*, both lacking β-GT and having higher ⍺-glucosylation levels than wild-type T4, T4Δ*β-gt* showed the lowest escaper frequency. In comparison, a 6-fold higher escaper frequency was observed with phage T4ΔP*_β-gt_*::*⍺-gt*, which could be related to its lower ⍺-glucosylation levels. The presence of β-GT, which consequently lowers ⍺-glucosylation levels, resulted in a further 2-fold higher frequency of escapers mutants in the wild-type T4 and T4ΔP*_⍺-gt_*::*β-gt*ΔP*_β-gt_*::*⍺-gt* phages (Figure 6A). Hence, decrease in the ⍺-glucosylation levels increases the escaper frequency until saturation is reached at low ⍺-glucosylation levels. These findings demonstrate that in the absence of type IV RM systems, the competition between β-GT and ⍺-GT increases the ability of T4 to escape Brig1 by lowering the ⍺-glucosylation levels.

**Figure 6:**
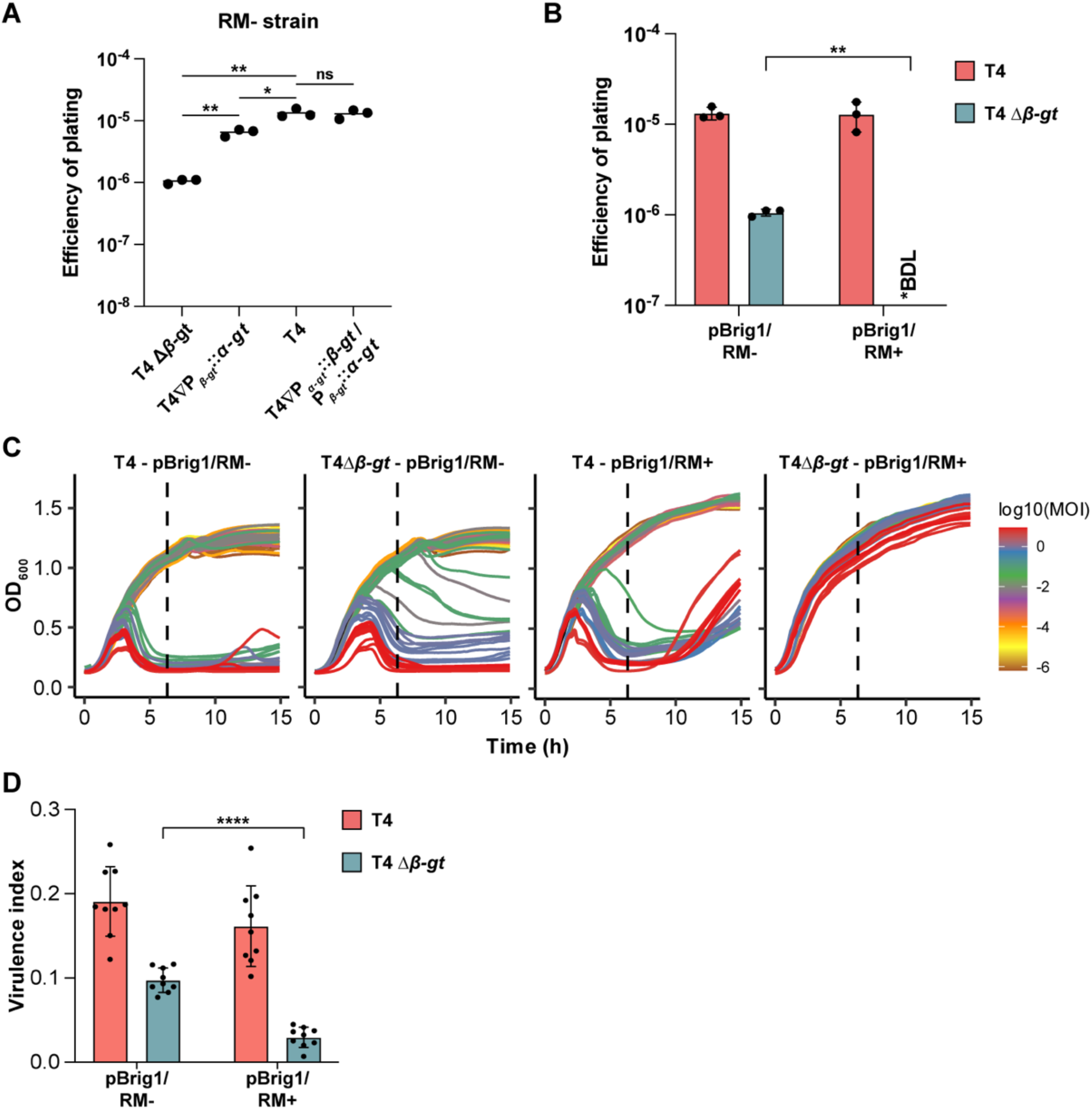
Role of β-GT in protection against Brig1. **A.** The frequency of Brig1 escapers, estimated as the efficiency of plating (EOP) of phages T41′*β-gt*, T4ΔP*_α-gt_*::*β-gt*, T4 and T4ΔP*_α-gt_*::*β-gt*/P*_β-gt_*::*α-gt* on *E. coli* DH10B carrying a *brig1* expressing plasmid. Data shown are mean of three biological replicates, the horizontal line represents the average value. A two-tailed unpaired t-test of the EOP values was used to calculate the *P* values; \*\**P* < 0.01, \**P* < 0.05, ns – not significant. **B.** Frequency of brig1 escapers (efficiency of plating of phages) T4 and T41′*β-gt* in *E. coli* strains DH10B (RM-) and MG1655 (RM+), both carrying a *brig1* expressing plasmid. The results are represented in terms of EOP. *BDL = Below Detection Limit. Data shown are mean of three biological replicates. A two-tailed unpaired t-test of the EOP values was used to calculate the *P* values; \*\**P* < 0.01. **C.** Growth curves of *E. coli* EC100 (RM-) and *E. coli* MG1655 (RM+) infected with phages T4 or T41′*β-gt* over a range of MOIs between 10^2^ to 10^−6^. Uninfected culture is represented in black. **D**. Virulence index of the respective phages, as calculated from the growth curves shown in (**C**), up to 7 hours post-infection. Data shown are mean of nine biological replicates, represented as mean value ± SD. A two-tailed unpaired t-test of the virulence index values was used to calculate the *P* values; ****P < 0.0001.

The formation of abasic sites during Brig1 activity results in escapers that are likely to contain single base pair deletions/insertions or substitution mutations, rather than large deletions typical of escapers generated through mini-homology mediated recombination ^45^. To confirm this, we analyzed the *⍺-gt* locus from escapers of Brig1 in comparison to escapers from CRISPR-Cas targeting of the same locus. Indeed, none of the 29 escapers from Brig1 targeting showed any difference in size, relative to the coding region from the wild-type phage, as opposed to the escapers of CRISPR-Cas targeting (Figure S14 and <u>Table S3</u>).

Next, we investigated the role of T4 β-GT in a scenario where the host encodes both DNA glycosylases and RM systems. In wild-type T4, Brig1 escaper frequency was independent of the presence of RM systems (Figure 6B). On the other hand, phage T4Δ*β-gt* was unable to generate escaper mutants in the presence of both RM and Brig1 (Figure 6B). To quantify the consequences of these results on a population level, we estimated the virulence index of phage T4 and T4Δ*β-gt* in hosts encoding Brig1 either in the absence or presence of RM systems. T4 virulence was independent of the presence of RM systems but phage T4Δ*β-gt* lacking β-GT was completely unable to propagate in the presence of both RM and Brig1 (Figure 6C, Figure S15 and Figure 6D). Based on these results we can conclude that in the presence of type IV RM systems, β-GT enables the survival of ⍺-GT inactivated Brig1 escapers by adequately glucosylating phage DNA.

Together, these results show that the antiphage activity of Brig1 and phage survival is partly dependent on the amount of ⍺-glucosylation within the phage DNA. We also show that the fitness of Brig1 escaper T4 mutants in a host strain encoding type I and type IV RM systems is entirely dependent on the phage encoding an alternative primary modification system such as *β-gt*. These results also demonstrate that in the presence of a single GT, Brig1 combined with RM systems can completely prevent emergence of phage escaper mutants.

## DISCUSSION

DNA hypermodification in phages is a general counter defense strategy against host defense systems whose mode of action is dependent on DNA sequence recognition, such as CRISPR-Cas and RM systems. Although it was known that bacteriophage T4 encodes two glucosyltransferases with similar functions, the mechanisms underlying their respective activities and the advantage of this functional redundancy had never been thoroughly investigated.

In phage T4, the cytosines are pre-replicatively hydroxymethylated and subsequently glucosylated to the fullest extent. We show that α-GT and β-GT, respectively, contribute to 66% and 33% of the total glucosylation, consistent with the prior estimates of 60% and 40%, respectively ^17,19^. Competition assay involving the two GTs demonstrated that, under *in vitro* conditions, β-GT exhibited greater enzymatic activity than α-GT, which contrasts with its lower contribution to glucosylation observed *in vivo* in wild-type T4. It has been previously hypothesized that the higher *in vivo* contribution of α-GT was due to a strong interaction between α-GT and the T4 replication machinery. Yet, under our tested *in vitro* conditions, neither the α-GT nor β-GT interacted with the T4 sliding clamp protein, T4 gp45, and metal-affinity purification using genomically tagged α-GT or β-GT revealed no interacting proteins. Rather, thanks to experiments of promoter switching in T4 genome, we show that the proportion of cytosines undergoing α- or β-glucosylation is significantly modulated by the transcriptional and translational activities associated with each GT, together with their enzymatic activity.

Despite strict size constraints on phage genomes, why do T4 and other closely related phages encode two primary GTs? We show that deleting either *gt* did not affect phage fitness, even in the presence of type IV RM systems targeting 5-hmC containing DNA. Moreover, the observation that, in the presence of α-GT alone, 80% of 5-hmC glucosylation fully protect phages against *E. coli* type IV systems indicates that this degree of glucosylation is sufficient for protection. Interestingly, we observed that the mutant phage T4ΔP*_β-gt_*::*α-gt*, with *α-gt* expressed from the *β-gt* promoter, makes smaller plaques and has a lower virulence index than other phages. The percentage of glucosylated 5-hmC was estimated to be 75%, a lower proportion of glucosylated bases as compared to the 80% glucosylation observed in single deletion genotypes. This suggest that 80% of glucosylation is the absolute minimal threshold necessary to protect against the *E. coli* McrBC/McrA RM system, an outcome that might be generalizable to other *E. coli* phages. In conclusion, in the wild-type phage, competition between the two enzymes ensures balanced distribution of the two types of linkages, but when expressed alone each enzyme utilizes either its expression strength or superior enzymatic activity to ensure adequate glucosylation and complete protection of phage DNA.

Here we also demonstrate that growth of phages encoding a single GT is entirely restricted in hosts encoding defense systems, such as Brig1, that target specific stereoisomers of 5-gmC, in addition to classical RM systems. On the contrary, phages encoding two different GTs can generate escape mutants at high frequency (1 × 10^−5^). Indeed, in the presence of a single GT, mutations inactivating α-GT leads to phage DNA targeting by type IV RM systems and are therefore lethal. By contrast, in the presence of two GTs with different stereospecificity, mutations in *α-gt* are offset by β-glucosylation, ensuring survival of escape mutants with inactivating mutations in *α-gt*.

Furthermore, among hosts encoding only the DNA glycosylase defense system, the presence of non-targeted GT(s) promotes a higher rate of escaper generation. Our results emphasize the benefits of GT diversity in phage genomes, and in the diversity of the types of modification on a more general level. These findings also underscore a notable drawback inherent to DNA glycosylase-based defense systems, such as Brig1, in that their activity invariably results in escaper phages in the absence of co-occurring defense systems. Analysis of Brig1 escapers across the α-gt locus showed that the mutants comprised only point mutations (single base pair insertions or deletions and substitutions), with none of the mutant’s showing larger deletions across or within *α-gt*, in contrast to what is observed in escapers from CRISPR-Cas systems ^45^. This constitutes an advantage for Brig1 escapers, as point mutations are reversible, offering the possibility to recover functional genes. Hence encoding two GTs whose products are stereoisomers confers an evolutionary advantage when faced with a stereoisomer specific host defense system.

The frequent association of β-glucosyl-HMC-α-GT homologs with both ⍺-GT and β-GT indicates that either the secondary GT does not differentiate between the two primary linkage types or that there are two distinct kinds of secondary GT: β-glucosyl-HMC-α-GT and β-glucosyl-HMC-β-GT. Furthermore, the presence of solitary secondary GT homologs, independent of both ⍺-GT and β-GT, in 288 out of 753 phage genomes indicates that these phages potentially encode novel primary transferases which forms the basis for additional glucosyl modifications. The not so limited prevalence of β-GT and arabinosylation clusters ^31^ among phages, along with the identification of diverse DNA hypermodification enzymes from metagenomic DNA ^46^, suggest that Brig1 is likely not the only DNA glycosylase based host anti-phage defense system. Similarly, it is also likely that the yet to be defined function of the secondary GTs lies in their ability to overcome DNA glycosylases that indiscriminately target primary GTs. To an extent, this has been demonstrated in Bas46 phage, where deletion of the secondary arabinosyltransferase Aat resulted in susceptibility to a plasmid encoded defense system^31^. Overall, only 6 out of the 753 phage genomes, including phage T4 and its close relatives, encoded both α-GT and β-GT. The limited co-occurrence of the two primary GTs might indicate that the adaptation of DNA glycosylase defense system by the hosts and the counter measure of encoding multiple hypermodification enzymes by phages is a relatively recent innovation.

Encoding multiple hypermodification enzymes such as GTs along with dCTPase, thymidylate synthase, IPI, IPII and IPIII is a great example of continuous antagonistic evolution of defense and counter defense systems. Phages hydroxymethylate their cytosines in response to sequence-specific targeting by the host type I RM system ^23^, which, in turn, makes them susceptible to the type IV McrBC RM system ^5,24^. This vulnerability is then countered by sugar modification, such as glucosylation or arabinosylation, of the cytosines ^17,18,31^. These modifications not only confer resistance to type IV RM systems but also enable the phage to evade targeting by certain types of CRISPR-Cas systems ^5–10,31^. Other type IV RM systems such as GmrSD are capable of indiscriminately targeting the DNA modified by the primary GTs, eliciting the evolution of phage encoded cognate inhibitors, IPI, IPII and IPIII ^25,26,28^. Subsequently, when hosts evolve stereoisomer-specific DNA glycosylases such as Brig1 ^27,32^, instead of evolving a cognate inhibitor, phage T4 has acquired a variant GT enzyme with different stereospecificity, *β-gt*, to diversify the distribution of the glycosidic linkages in its genome (Figure 7).

**Figure 7:**
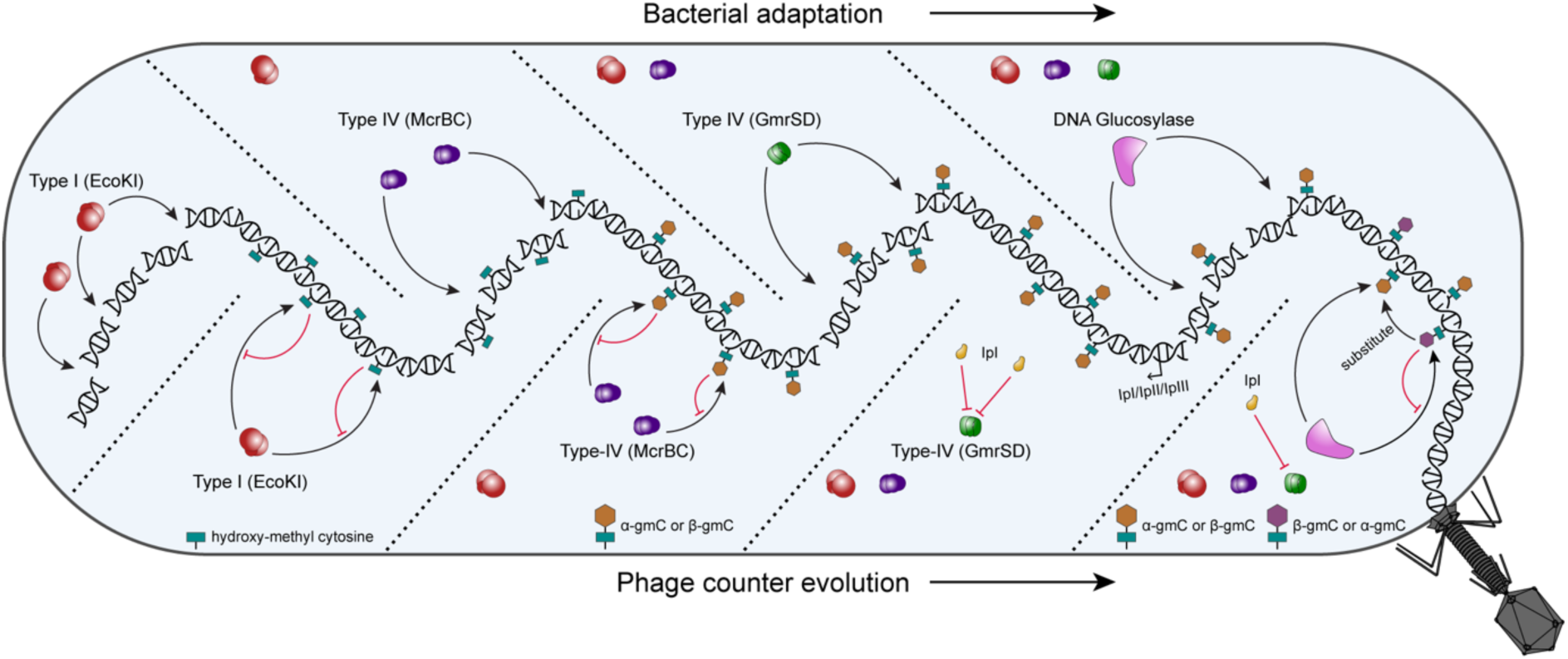
Model of T4 counter evolution to overcome host defense systems. From left to right, upon evolution of the type I RM systems, phages likely adapted a host enzyme to hydroxy methylate the cytosine base in their genome. In response, host evolved the Type IV McrBC RM system, which was in turn countered by hypermodification of the 5-hmC in the phage genome. The hypermodification was in turn countered by Type IV GmrSD RM systems, which are suppressed by the phage protein IpI. Following this, evolution of DNA glycosylases such as Brig1, specific to α-glucosylation, are countered by acquisition of β-GT enabling viability of Brig1 escapers.

## Supporting information

Identification and analysis of GT homologs in phage genomes from the RefSeq and non-redundant (nr) NCBI GenBank database

List of bacterial and phage strains, plasmids, and oligonucleotides used in this study

## DATA AVAILABILITY

Supporting data linked to the findings in this study can be found in the main article or the supplementary materials. Further data will be made available from the corresponding author upon request.

## ACKNOWLEDGEMENTS

The authors thank all members of the MuSE lab (Micalis), INRAE, Jouy-en-Josas for the insightful discussions; Safir Hien Phan (University of Copenhagen, Denmark) for her technical support; Luciano A. Marraffini (The Rockefeller University, USA) for providing the pBrig1 plasmid and *E. coli* EC100 strain. We would also like to thank the BSRC Mass Spectrometry & Proteomics Facility, University of St. Andrews and PAPPSO, INRAE for their help with proteomic analysis. The authors acknowledge a DIM1Health 2019 grant from the Région Île-de-France to the project EpiK for the LC-MS equipment. This work was supported by the ANR grant MUMI [ANR-20-CE12-0008-02] to M.D.P. and the Novo Nordisk Fonden Postdoctoral Fellowship in Bioscience and Basic Biomedicine Grant [NNF21OC0067491] to Y.B.-C.

## AUTHOR CONTRIBUTIONS

Conceptualization, Y.B.-C. and M.D.P.; investigation, Y.B.-C., L.R.-C., F.B. and A.L.I.W.; formal analysis, L.R.-C., M.K., F.B., Y.B.-C. and F.L.; validation, L.R.-C., F.B. and A.L.I.W.; visualization Y.B.-C., L.R.-C. and M.K.; funding acquisition, Y.B.-C., M.D.P. and M.H.; supervision, Y.B.-C., M.D.P. and M.H.; writing – original draft, Y.B.-C., M.K., L.R.-C., F.B. and M.D.P; writing – review and editing, M.D.P., F.L., M.K., Y.B.-C. and M.H.

## METHODS

### Growth conditions, PCR, plasmids and strains

*Escherichia coli* MG1655 was utilized for the growth curve assays, involving the estimation of virulence index, and *E. coli* DH10B was utilized in genome editing experiments and plaque assays. All *E. coli* cultures were grown in Luria-Bertani (LB) medium, incubated at 37°C with shaking at 200 rpm.

Oligonucleotides and PCR fragments were fused together either by Overlap Extension-PCR or by Gibson Assembly (NEB #E2611S) ^47^ and cloned into the plasmid pBAD24. In case of the positional swapping between *α-gt* and *β-gt*, the coding sequence of β-GT was first replaced with a pseudo-gene whose sequence includes appropriate protospacers for targeting by the CRISPR-Cas13b system. A pBAD24 based plasmid was constructed with the coding sequence of α-GT fused to the homologous arms that correspond to the sequences flanking the coding sequence of β-GT. Similarly, a plasmid containing the coding sequence of β-GT flanked by the upstream and downstream sequence of α-GT was also constructed. A similar strategy was used to insert glucosyltransferases with in-frame histidine coding sequence.

Hydroxymethylated deoxycytosines (hmdC) containing DNA fragment was prepared by PCR amplification of DNA fragment using the primers T4 DNApol For and T4 DNApol Rev with T4 DNA as template. In the PCR preparation, the dNTP mixture was substituted with an alternative mixture that was prepared by mixing the individual nucleotides, dATP, dGTP and dTTP along with 5-Hydroxymethyl-dCTP (Jena Bioscience, #NU-932S).

Spacers corresponding to target protospacers in *α-gt*, *β-gt* and pseudo-gene were introduced into the plasmid pBZCas13b (Addgene #103986), encoding Cas13b, as described earlier. Briefly, overlapping oligonucleotides with 5’ and 3’ extensions were pooled together at 1 μM final concentration, along with PCR reaction buffer. The mixture was incubated at 95°C for 10 minutes and gradually cooled to room temperature to facilitate annealing. The annealed oligonucleotides were ligated into the pBZCas13b vector linearized with the BsaI-HF (NEB #R3733S).

For heterologous expression of individual proteins, the coding sequences were cloned into the pET-28a(+) vector while retaining an in-frame C-terminal histidine sequence (in the case of *α-gt* and *β-gt*) or without an in-frame histidine coding sequence (T4 *gp45*).

All primers, plasmids and strains used in this study are listed in Table S4.

### Bioinformatic analysis

Homologs of the three glucosyltransferases, NP_049673.1 (α-GT, pfam11440), NP_049658.1 (β-GT, pfam09198) and YP_010067197.1 (β-glucosyl-HMC-α-GT, pfam20691), encoded among viruses (taxid = 10239), in the RefSeq database were identified using Position Specific Iterative BLAST (PSI-BLAST) ^48^, with an expect threshold = 0.005, over three iterations. The protein ids of the homologs of all three glucosyltransferases were combined and organized according to their respective phage accession numbers, which was then utilized to visualize their distribution. A similar analysis was carried out against all viruses in the non-redundant (nr) NCBI GenBank database.

VICTOR (Virus Classification and Tree Building Online Resource) ^49^ was used to construct the phage phylogenetic tree based on phages retrieved from the RefSeq database that encode at least one GT and belong to the genus *Tequatrovirus*. Representative phages from diverse genera that also encode at least one GT were also included. Pairwise comparisons of the phage nucleotide sequences were conducted using the Genome-BLAST Distance Phylogeny method (including 100 pseudo-bootstrap replicates each) with the D0 formula for nucleotides ^50^. The tree was visualized with ggtree ^51^. To construct a phylogenetic tree of T6 β-glucosyl-HMC-α-GT, protein homologs were retrieved from RefSeq database during the earlier PSI-BLAST and submitted to the phylogenetic analysis tool NGphylogeny.fr with default settings ^52^. The Newick tree generated by PhyML was visualized on iTOL ^53^.

For comparison of the six phage genomes, the complete nucleotide genome sequences of *Escherichia* phage 236Ecol005PP (PP434423.1), *Escherichia* phage FL12 (PP400786.1), *Escherichia* phage Pu-Krd-SF1 (PQ9395.1), *Escherichia* phage T4 (NC_000866.4), *Enterobacteria* phage RB55 (KM607002.1) and *Escherichia* phage vB-Eco-KMB26 (OR539218.1) were downloaded in fasta format and combined into a single file. The nucleotide sequences were then analyzed using VIRIDIC WEB ^54^, to calculate the nucleotide similarity between each of the six genomes, with default BLASTN parameters (‘-word_size 7 -reward 2 -penalty −3 - gapopen 5 -gapextend2’).

### Three-dimensional structure prediction and analysis

PDB identifiers for representatives of different GT families were retrieved from the CAZy database ^41^. All viral and cellular protein structures were downloaded from the PDB database ^55^. Structural models of phage proteins for which experimentally determined structures were not available were either downloaded from the BFVD database ^56^ or modelled using AlphaFold3 ^57^. Structural similarities between cellular and viral proteins were evaluated based on the DALI Z score, which is a measure of the quality of the structural alignment. Z scores above 2, i.e., two SDs above expected, are usually considered significant ^58^. Structural similarity matrix from all-against-all structure comparisons as well as corresponding dendrograms were obtained using the latest release of the DALI server ^40^. Structures were visualized using the University of California, San Francisco (UCSF) ChimeraX v1.9 ^59^.

### Protein purification

200 ml cultures of *E. coli* BL21(DE3) strains (Novagen) carrying pET-28a(+)_*α-gtchis* or pET-28a(+)_*β-gtchis* were grown in LB medium at 37°C, 200rpm until OD_600_ = 0.6-0.8 was reached. Cultures were then incubated at 30°C for 3 hours and protein expression induced by the addition of IPTG to a final concentration 0.5 mM. Cells were collected by centrifugation at 6300 x g for 10 minutes at 16°C. The pellets were resuspended in lysis buffer (25 mM Tris-Cl, 200 mM NaCl, 5% glycerol and 30 mM Imidazole, pH = 7.5) and stored at −16°C until further processing. The thawed mixture was lysed by homogenization (STANSTED model SPCH-10 from homogenizing systems, UK) and sonication (30 cycles, 3 seconds ON and 3 seconds OFF). The cell debris was removed by centrifugation at 12000 x g for 30 minutes at 4°C and the supernatant was filtered with a 0.45 μM filter. Proteins were bound to histrap columns (HisTrap™ High Performance column, Cytiva), equilibrated with the lysis buffer. After washing with 40 column volumes of the lysis buffer including 30 mM Imidazole, the proteins were eluted in a buffer including 500 mM Imidazole. The eluted fractions, containing the protein of interest, were pooled together and concentrated using Pierce™ Protein Concentrators PES 10K MWCO (ThermoFisher SCIENTIFIC, #88516). The concentrated fractions were loaded on a Superdex 200 Increase 10/300 GL column (Cytiva) equilibrated in 25 mM Tris-Cl, 200 mM NaCl, 5% glycerol, pH = 7.5 (SEC buffer).

### Analysis of α-GT-gp45 and β-GT-gp45 interactions, *in vitro*

For the analysis of *in vitro* protein-protein interaction, induction of pET-28a(+)_*gp45*, along with pET-28a(+)_*α-gtchis* or pET-28a(+)_*β-gtchis* was performed as described above. The filtered supernatants containing tag-free gp45 was split into two fractions and mixed with supernatants containing α-GT chis or β-GT chis. The supernatants were then combined with Ni-NTA Agarose (Qiagen, #30210), equilibrated with the lysis buffer. The mixtures were incubated at 4°C for 60 minutes, washed with lysis buffer several times and eluted with a buffer containing 500 mM Imidazole. Filtered supernatants of the individual proteins were also treated similarly and purified, as controls.

### *In vivo* pull-down with endogenously tagged α-GT and β-GT

The pull-down assay was carried out as described previously ^20^, with minor modifications. *E. coli* MG1655 cultures in exponential phase (OD_600_ = 0.5-0.6) were infected with mutant phages, that encoded a histidine tagged α-GT or β-GT, at an MOI of 3. The infected culture was then incubated for 15 min at 37 °C, chilled on ice and centrifuged at 6000 xg for 10 min at 4 °C. Pellets were conserved at −20 °C until further use. Pellets were resuspended in a lysis solution (50 mM Tris-HCl pH = 7.5, 250 mM NaCl) mixed with cOmplete™, Mini, EDTA-free Protease Inhibitor Cocktail (Sigma #11836170001) and sonicated at 200 W 5 cycles, 10 sec on, 10 sec off per cycle (Sonics cat. VC505). 1 µL of Turbo DNase I (Invitrogen #AM2238) was added to the lysate, which was then centrifuged at 10 000 xg for 10 min. Supernatants were filtered through 0.20 µm PVDF (Fisher # 097203) and incubated overnight at 4 °C in contact with Ni resin (NEB #S1428S). The protein bound resins were washed thrice with 20 mM Tris-HCl pH = 7.5, 150 mM NaCl and 30 mM Imidazole. Finally, the bound proteins were eluted with 20 mM Tris-HCl pH = 7.5, 150 mM NaCl and 500 mM Imidazole.

### Competition assay

Substrate DNA containing 5-hmdC was amplified using the primers T4 DNApol For and T4 DNApol Rev as described earlier. The glucosylation reaction was carried out in NEB buffer 4 and included 740 ng of DNA (equivalent to 400 pmol of 5-hmdC, 16 pmol/μl final concentration), Uridine-5’-diphosphoglucose disodium salt (MP Biomedicals, LLC, Cat no. 101208) to a final concentration of 2 μM, along with the appropriate amounts of GT, to a final concentration of 1.5 pmol/μl. The reactions were incubated at 37°C for 60 minutes, following which the products were directly digested using the Nucleoside digestion mix and quantified by LC-MS/MS (described below).

### Phage preparation and plaque assay

Phage stocks were prepared as lysates by infection of *E. coli* DH10B cultures in exponential phase (OD_600_ 0.1-0.2) in LB with 10 mM MgSO_4_ and 5 mM CaCl_2_. After the infected cultures reached an OD_600_ lower than 0.1, they were centrifuged and filtered. The stocks were titrated and stored at 4 °C until use.

For plaque assay, preheated top agar (0.4% LB-agar) was mixed with 100 μl of overnight culture and phage preparation. The mixture was then poured onto preheated LB agar plates and incubated overnight at 37 °C. The plaques were counted and the plaque-forming unit per ml (PFU/ml) was calculated.

### Genome editing of phage T4

Insertions and deletions in bacteriophage T4 were performed as described earlier ^43^. Briefly, overnight cultures of the recombinant strains, E. coli carrying plasmids constructed for deletions or insertions, were diluted and incubated at 37°C until an OD_600_ between 0.15-0.25 was reached. The cultures were then infected with phage T4 preparations at an MOI of 0.05 and incubated at 37°C until complete lysis was observed (1-2 hours post infection). Serial dilutions of the supernatants were spotted on LB plates with the top layer containing the selection strain, *E. coli* DH10B with spacer containing pBZCas13b plasmid, to target the non-recombinant phage T4. The overnight incubated plates were then screened for the presence of the appropriate mutation by PCR.

Although replacement of the ⍺-GT coding sequence with *β-gt*, resulting in a recombinant phage with two copies of *β-gt*, is possible to construct using CRISPR-Cas based genome editing, subsequent replacement of the native β-GT coding sequence with *⍺-gt* becomes unachievable, as the CRISPR spacer would target both *β-gt* copies. Instead, we developed an alternative strategy in which we replaced the β-GT coding sequence with a pseudo-open reading frame (pseudo-ORF) in wild-type T4 phage, to construct phage T4 LySE. The pseudo-ORF carried three protospacers, designed for efficient targeting by the CRISPR-Cas13b system (Figure S8A). Utilizing spacers designed for targeting ⍺-gt transcripts, we introduced the β-GT coding sequence downstream of the *⍺-gt* promoter (referred to as T4ΔP*_⍺-gt_*::*β-gt*). Subsequently, this recombinant phage was further engineered to replace the pseudo-ORF with the ⍺-GT coding sequence downstream of the *dCMP*/*β-gt* promoter (referred to as T4ΔP*_⍺-gt_*::*β-gt* P*_β-gt_*::*⍺-gt*) (Figure 4B, Figure S8B). Another phage with single replacements was also constructed, referred to as T4ΔP*_β-gt_*::*⍺-gt*, in addition to T4ΔP*_⍺-gt_*::*β-gt*.

### Quantification of nucleosides by LC-MS/MS

Total phage genomic DNA was extracted as described earlier ^43^. Phage suspension was mixed with PEG and NaCl to a final concentration of 10% and 0.5 M respectively and incubated overnight at 4°C. Phages were pelleted by centrifugation at 12,000 rpm for 30 minutes at 4°C and resuspended in SM buffer (50 mM Tris-HCl pH 7.5, 100 mM NaCl, 8 mM MgSO_4_). Total DNA was extracted using Phenol-Chloroform followed by ethanol precipitation. 1 μg of total DNA was digested in 20 μl reaction volume using the NEB Nucleoside digestion mix (#M0649S) according to the manufacturer’s instructions.

Analysis of global levels of dC, 5mdC, 5hmdC, 5(α)-gmdC and 5(β)-gmdC were performed on a Q Exactive Orbitrap Mass Spectrometer (ThermoFisher Scientific) equipped with an electrospray ionization source (H-ESI II Probe) coupled with an Ultimate 3000 RS HPLC (ThermoFisher Scientific). Digested DNA was injected into ThermoFisher Hypersil GOLD aQ chromatography column (100 mm * 2.1 mm, 1.9 μm particle size) heated at 30°C. The flow rate was set at 0.3 mL/min and run with an isocratic eluent of 1% ACN in water with 0.1% formic acid for 10 min. Parent ions were fragmented in positive ion mode with 10% normalized collision energy in parallel-reaction monitoring (PRM) mode. MS2 resolution was 17,500 with an AGC target of 2e5, a maximum injection time of 50ms, and an isolation window of 1.0 m/z. The inclusion list contained the following masses: dC (228.1), 5mdC (242.1), 5hmdC (258.1) and 5(α)-gmdC/5(β)-gmdC (420.2). Extracted ion chromatograms of base fragments (±5 ppm) were used for detection and quantification (112.0506 Da for dC; 126.0662 Da for 5mdC; 142.0609 Da for 5hmdC; 304.1133 Da for 5(α)-gmdC/5(β)-gmdC). 5(α)-gmdC and 5(β)-gmdC were distinguished by their retention times. Calibration curves were previously generated using synthetic standards (Clinisciences, France) in the ranges and 0.01–1 pmole.

### Virulence index

Virulence index was calculated as previously reported ^44^. Briefly, 300 µL of overnight culture of *E. coli* MG1655, encoding McrBC/McrA RM systems targeting 5-hmC DNA, was used to inoculate 30 mL of LB, which was then incubated for 1 hour at 37 °C while being agitated at 200 rpm. Once the cultures reached OD_600_ between 0.15-0.25, 100 µL was distributed in a 96-well plate. In parallel, pre-titrated phage lysates were serially diluted and 100 µL of each dilution was added in to each well, covering a range of multiplicity of infections (MOIs) from 10^2^ to 10^−6^. The Optical Density (OD_600_) of the cultures was measured every five minutes over 16 hours to obtain bacterial reduction curves.

The limit of integration was set at 7 hours post infection, since beyond this time point re-growth of phage-resistant mutants was observed. Area under the curve (AUC) of the bacterial reduction ones were obtained through the function MESS::auc depicted as local virulence curves (AUC vs log_10_(MOI)). The virulence index of each phage was obtained by integrating the virulence curves and normalizing. Virulence index values were compared by Kruskal Wallis Pairwise comparison and Dunn’s all-pairs test.

### Efficiency of plating assay

Infectivity of phages were estimated through efficiency of plating (EOP) assays. Ten-fold serial dilutions were prepared from a single stock of the phages. 10 μl of phage dilution and 100 μl of overnight bacterial culture was mixed with preheated top agar (0.4% LB-agar) and poured on LB agar plates. After overnight incubation at 37 °C, the number of plaques was counted and the PFU/ml was determined for both the test and control strains. EOP was calculated as the ratio of their PFU/ml values.

## SUPPLEMENTAL FIGURES

**Figure S1:**
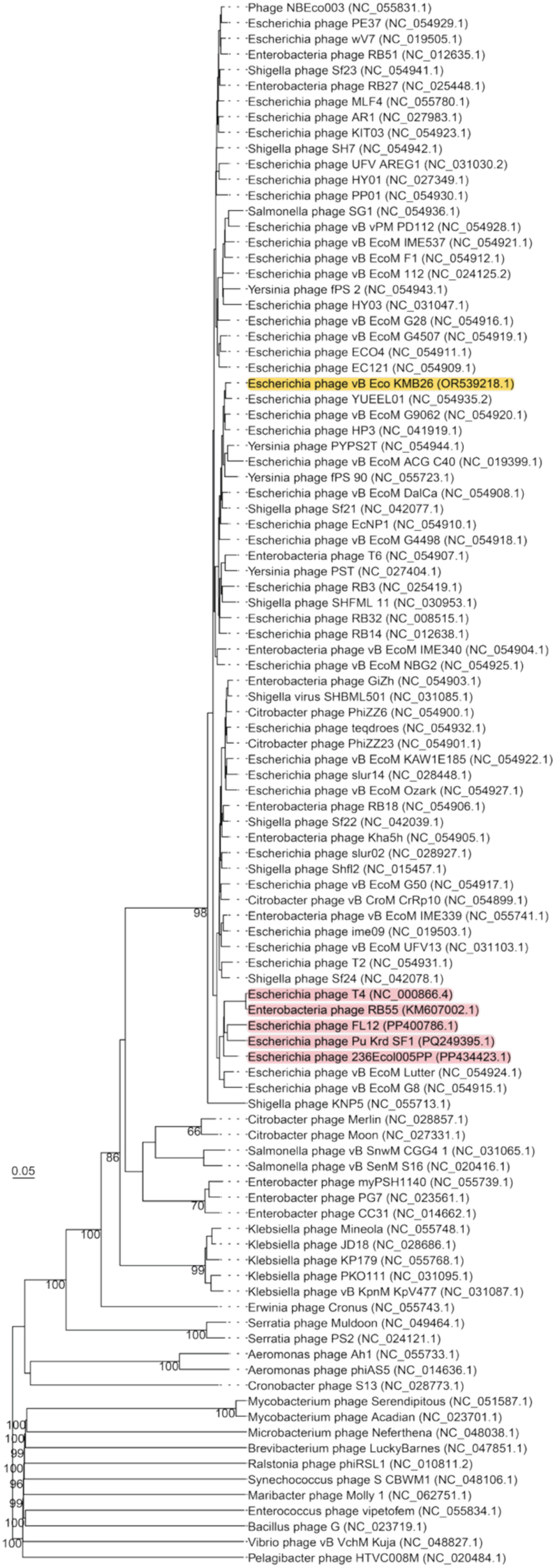
Phylogenetic tree based on the whole-genome comparison of phages encoding GTs. The phages highlighted in red encode both primary GTs and the phage highlighted in yellow encodes all three GTs. The Genome-BLAST Distance Phylogeny pseudo-bootstrap support values from 100 replicates are shown above the branches. The tree was generated using the VICTOR web service (https://ggdc.dsmz.de/victor.php) ^49^, with branch lengths scaled according to the nucleotide distance formula (D0).

**Figure S2:**
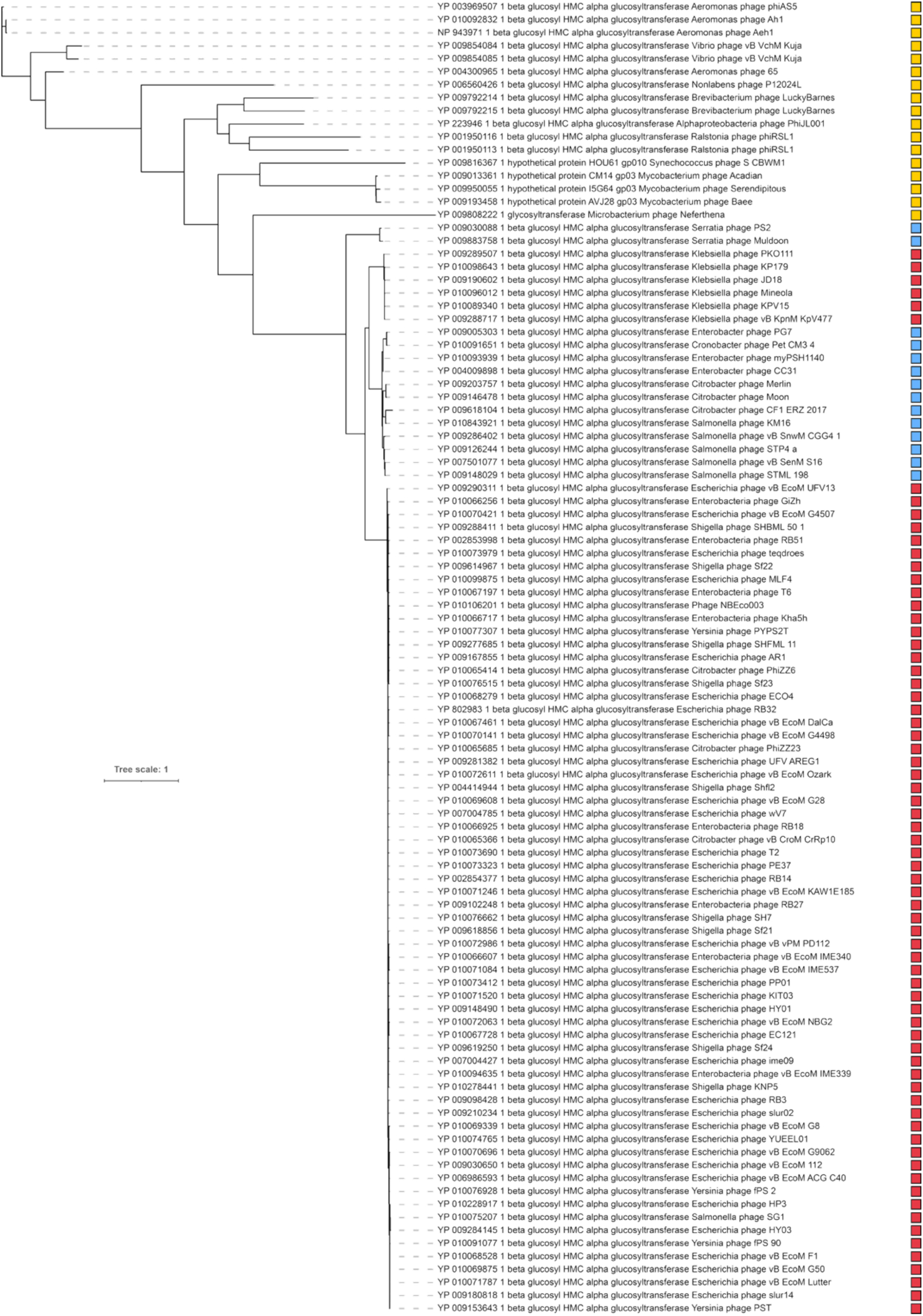
Maximum likelihood phylogeny of the secondary glucosyltransferase (β-glucosyl-HMC-α/β-GT). The box adjacent to each leaf label indicates co-occurrence with either α-GT (red) or β-GT (blue), or lack of association with a primary GT (yellow). The tree was generated using the NGphylogeny.fr ^52^ web service.

**Figure S3:**
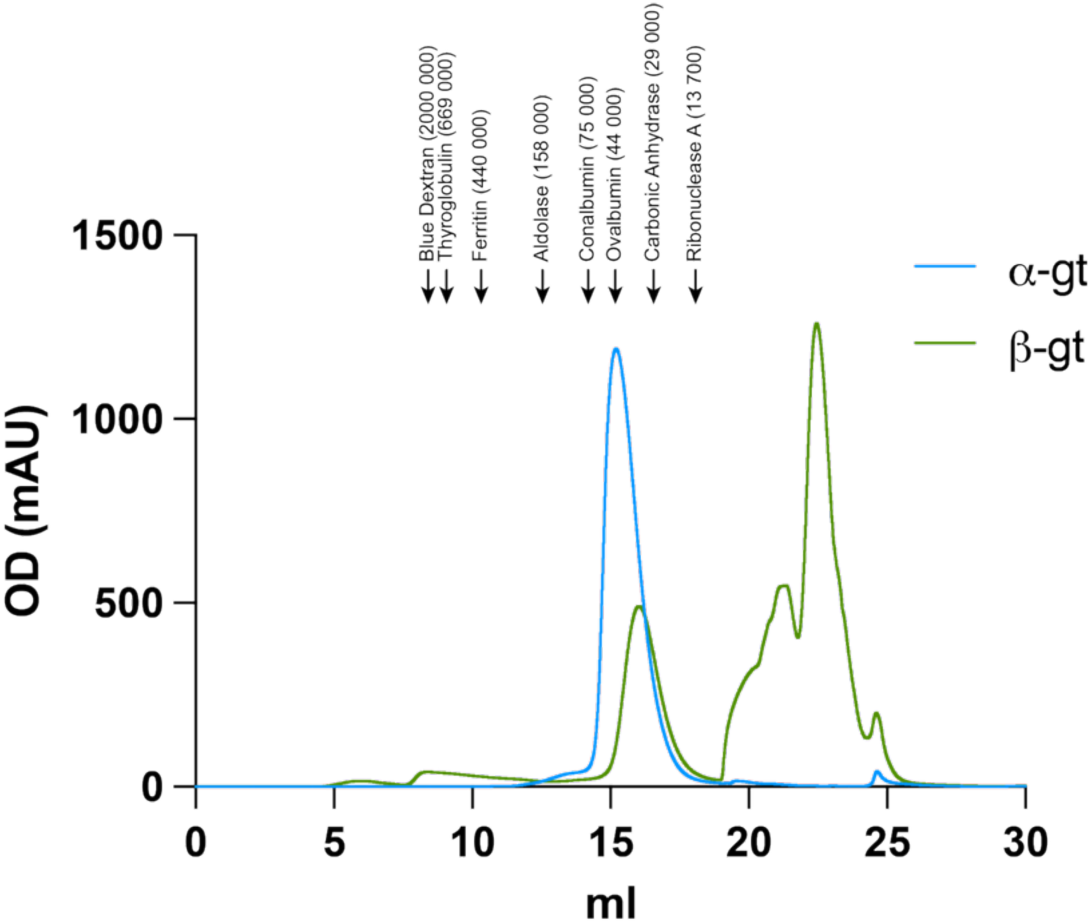
Size-Exclusion Chromatography of C-terminally histidine tagged ⍺-GT and β-GT. Elution profiles of histidine tagged α-GT (blue) and β-GT (green) obtained upon separation of the proteins on a Superdex 200 Increase 10/300 GL column. The absorbance was monitored at 280 nm. The peak elution position of each standard protein is indicated as an arrow, along with their theoretical molecular weights (Da) in brackets. The molecular weights of α-GT and β-GT, based on their peak elution volumes, were estimated to be 48.36 kDa and 33.61 kDa, respectively.

**Figure S4:**
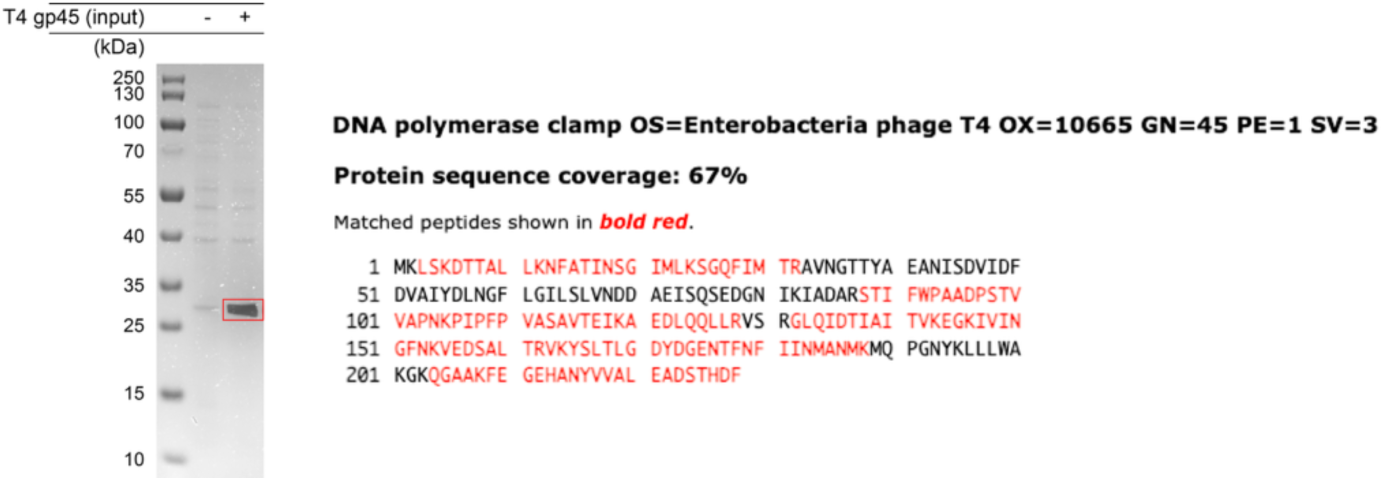
Validation of un-tagged T4 gp45 expression in *E. coli* BL21(DE3). Lane 1 and lane 2 correspond to the total cell extract of *E. coli* BL21(DE3) with un-tagged T4 gp45, before and after induction with IPTG. The induced protein (intense band) corresponding to the size of the un-tagged sliding clamp (T4 gp45, 24858.32 Da) monomer, was excised from the gel (shown inside the red box) and the protein’s identity confirmed by LC-MS/MS (right panel).

**Figure S5:**
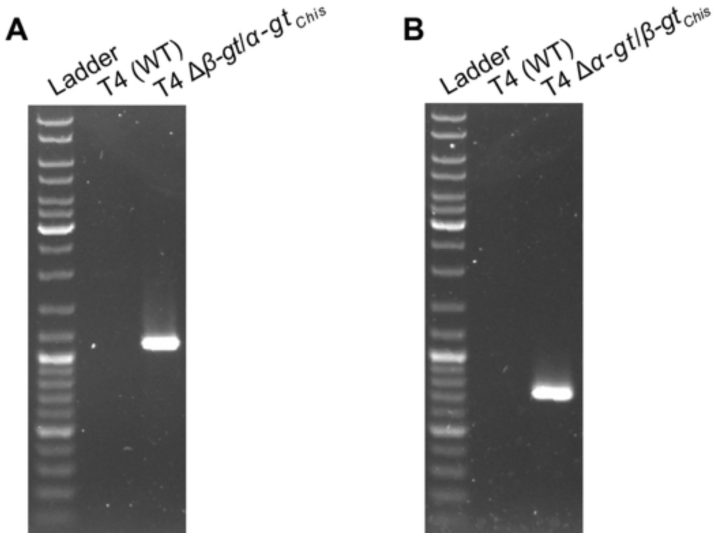
Sequence confirmation of the histidine-tag at the C-terminus of the GTs in the phage mutants T4 Δ*β-gt/⍺-gt_Chis_* and T4 Δ*⍺-gt/β-gt_Chis_*. **A.** PCR confirmation of the histidine tag coding sequence downstream of *α-gt* in the phage T4 Δ*β-gt/⍺-gt_Chis_* using primers T4_*⍺-gt*_chk_in_rev and Histag-C_chk_rev (listed in <u>Table S4</u>). T4 wild-type was used as control. **B.** PCR confirmation of the histidine tag coding sequence downstream of *β-gt* in the phage T4 Δ*⍺-gt/β-gt_Chis_* using primers T4_*β-gt*_chk_in_rev and Histag-C_chk_rev (listed in <u>Table S4</u>). Wild-type T4 was used as control.

**Figure S6:**
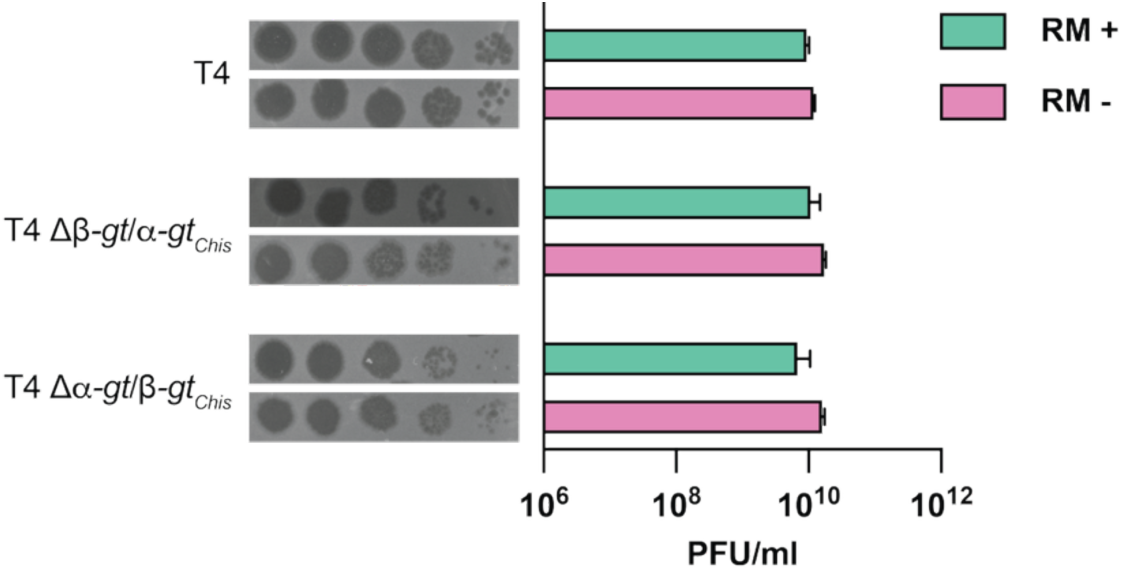
Estimation of the infectivity of phages T4 Δ*β-gt/⍺-gt_Chis_* and T4 Δ*⍺-gt/β-gt_Chis_* in the presence of RM systems. Phage spot assays demonstrating the infectivity of the phage mutants T4 Δ*β-gt*/*⍺-gt_Chis_* and T4 Δ*⍺-gt*/*β-gt_Chis_* in comparison to the wild-type phage. The spot assays were performed on *E. coli* MG1655 (RM +) and *E. coli* DH10B (RM −). The phage titers (PFU/ml), estimated from the spot assay, are from three biological replicates, represented as mean ± SD.

**Figure S7:**
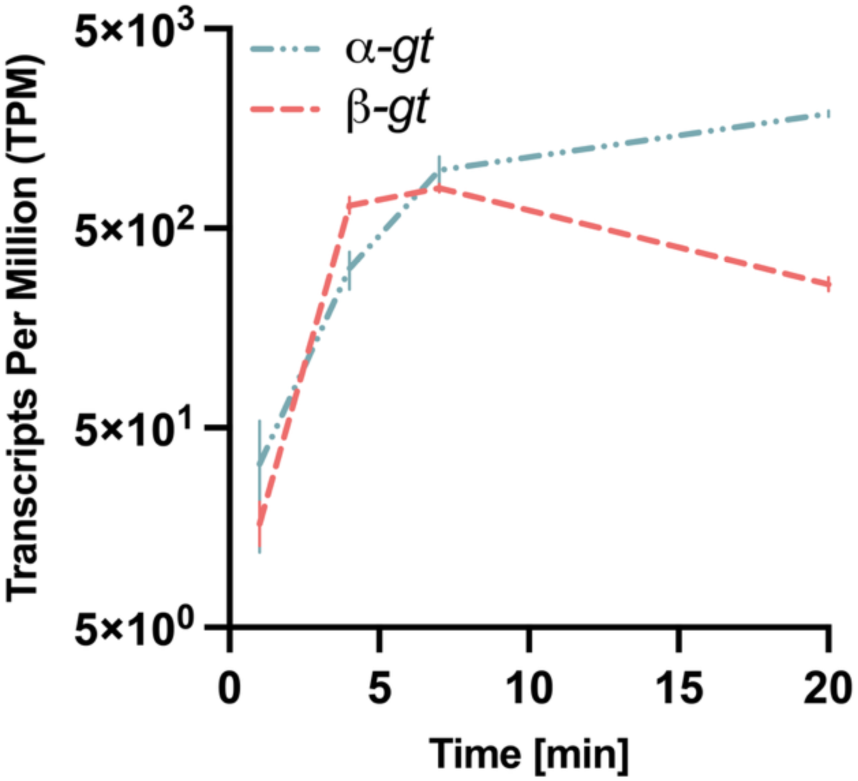
Transcript levels of *α-gt* and *β-gt* during the T4 infection cycle. Average of the three replicates, each normalized with the total reads corresponding to phage transcripts, was plotted against time post infection. Raw data was obtained from Wolfram-Schauerte and Pozhydaieva et al., 2022 ^42^.

**Figure S8:**
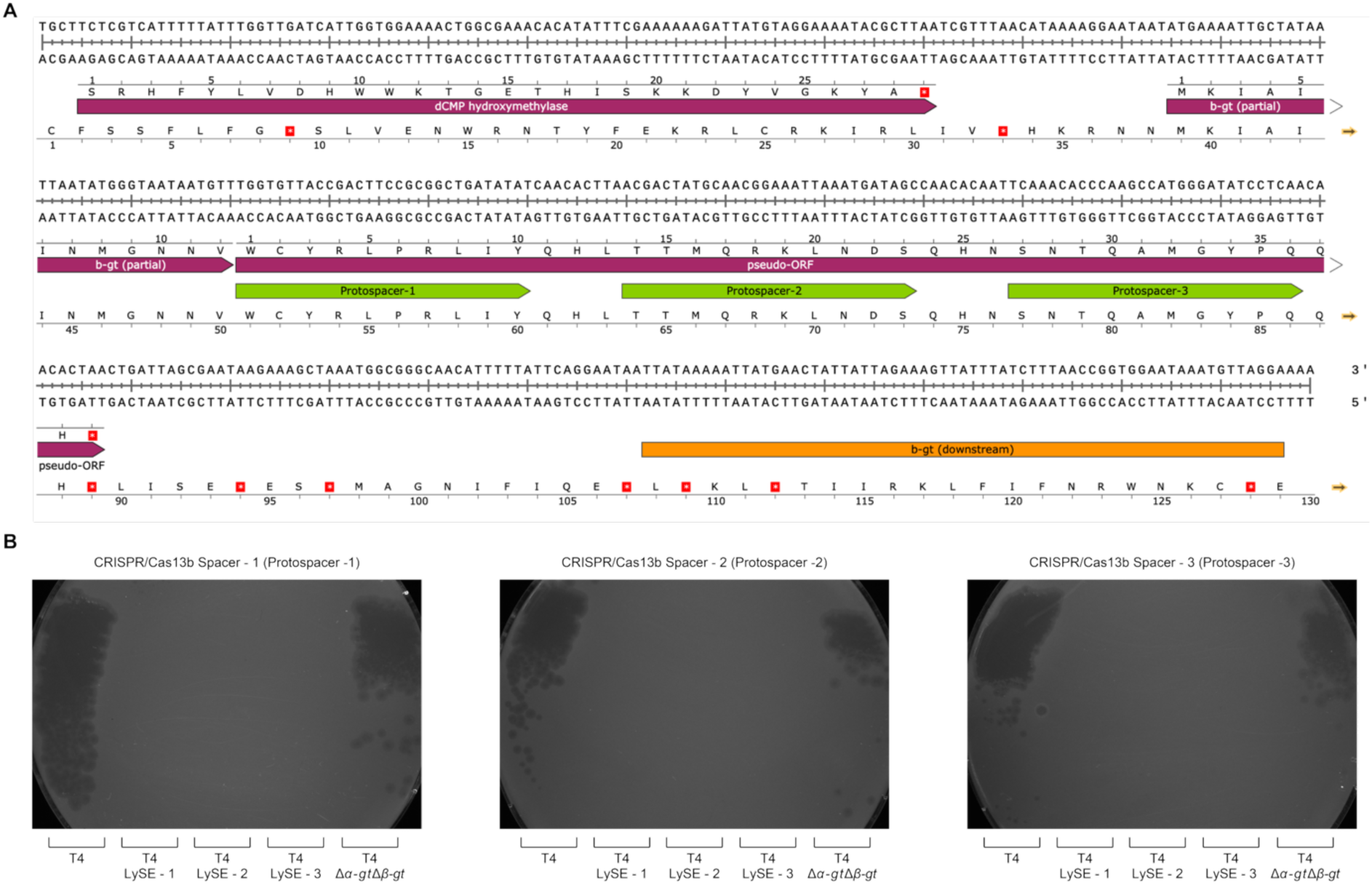
Genome editing strategy for swapping genomic locations of *α-gt* and *β-gt*. **A.** The coding region including the pseudo-ORF, in-frame, with the partial *β-gt* sequence is shown. The protospacers for Cas13b targeting are shown as arrows (in green). The upstream (*gp42* and partial *β-gt*) and downstream (*β-gt* downstream) sequences, necessary for recombination into the wild-type T4, are shown in purple and orange respectively. **B.** Susceptibility of the phage T4 LySE (3 clonal copies) to Cas13b is shown. Left panel: Cas13b spacer complimentary to protospacer–1, middle panel: spacer complimentary to protospacer–2 and right panel: spacer complimentary to protospacer–3. T4 wild-type and T4Δ*α-gt*Δ*β-gt*, both lacking the pseudo-ORF, are shown as controls.

**Figure S9:**
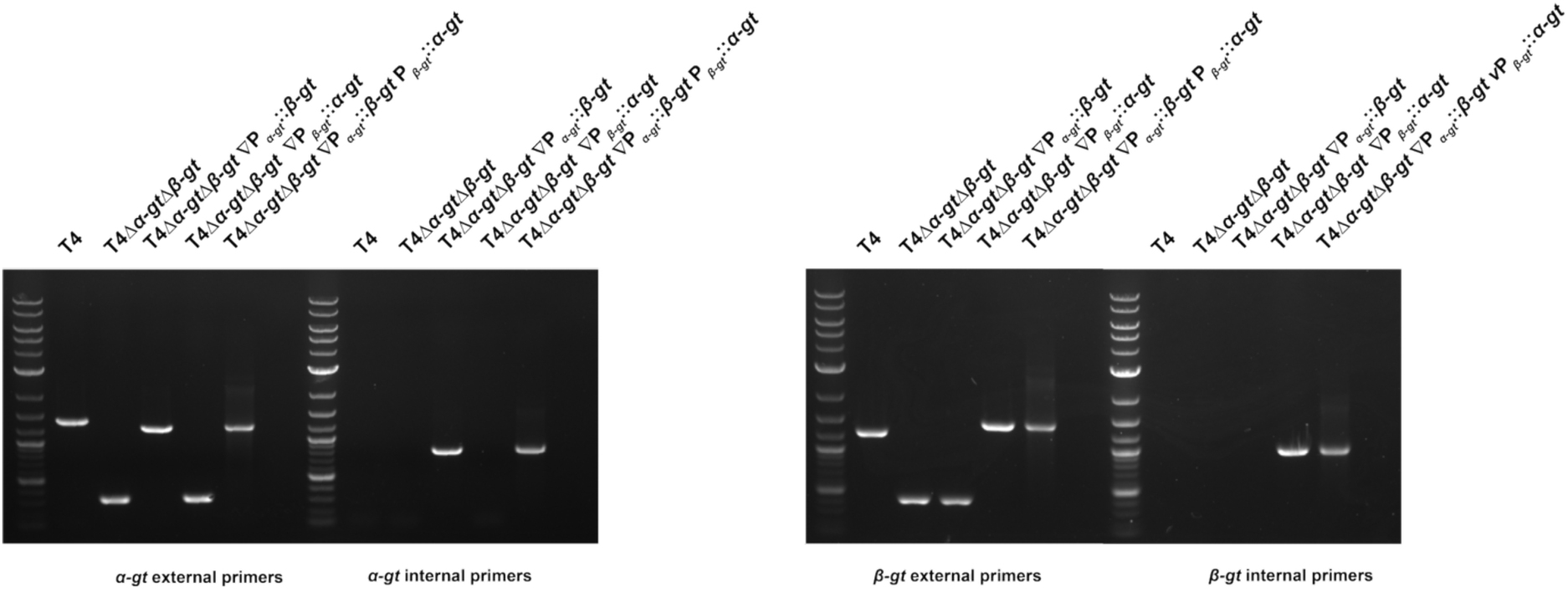
PCR confirmation of the *⍺-gt* and *β-gt* genomic location swapping in different phage mutants. The positional swapping of *α-gt* and *β-gt*, along with their presence and absence, in the phage mutants T4 ΔP*_⍺-gt_*::*β-gt*, T4 ΔP*_β-gt_*::*⍺-gt* and T4 ΔP*_⍺-gt_*::*β-gt* P*_β-gt_*::*⍺-gt* was confirmed with internal primers and external primers for *⍺-gt* and *β-gt* (listed in <u>Table S4</u>). Phage T4 was used as PCR control.

**Figure S10:**
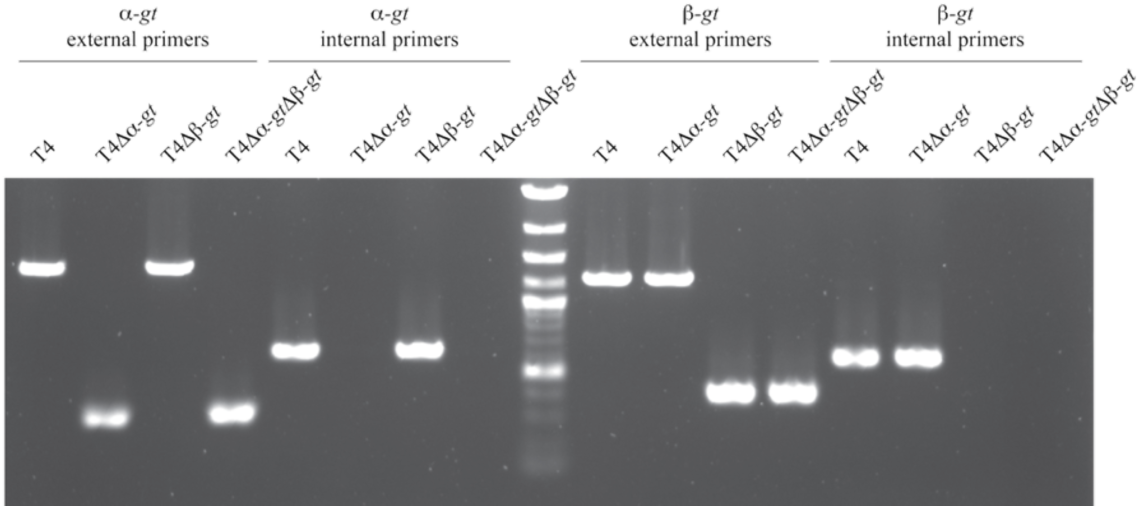
PCR confirmation of the desired deletion in T4Δ*α-gt*Δ*β-gt*. The deletion of *α-gt* and *β-gt* in the mutant phage T4Δ*α-gt* Δ*β-gt* is confirmed with internal primers and external primers (listed in <u>Table S4</u>) for *⍺-gt* and *β-gt*. Mutant phages T4Δ*⍺-gt* and T4Δ*β-gt*, constructed earlier ^43^, along with the phage T4 were used as PCR controls.

**Figure S11:**
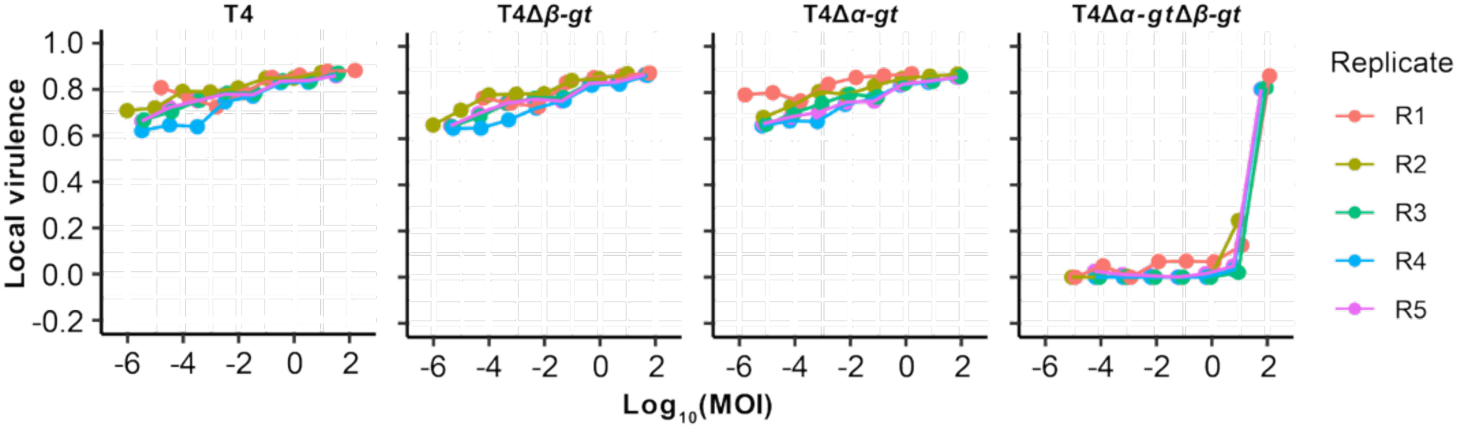
Local virulence corresponding to phages T4, T41′*α-gt*, T41′*β-gt* and T41′*α-gt*1′*β-gt* upon infection in *E. coli* MG1655. The local virulence for each MOI was calculated from the corresponding growth curve in Figure 5A.

**Figure S12:**
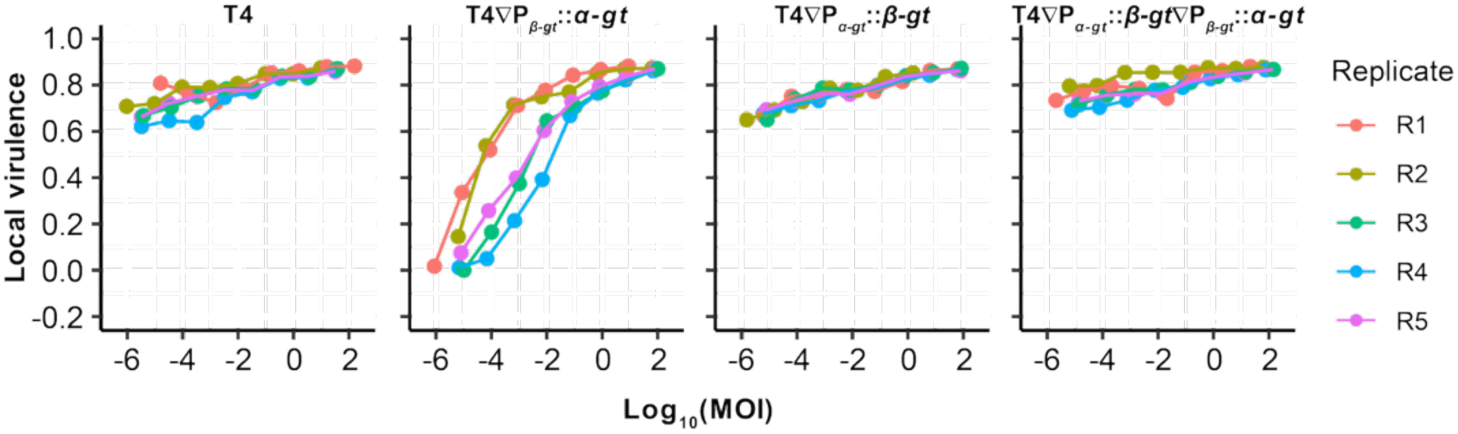
Local virulence of phages T4 ΔP*_α-gt_*::*β-gt*, T4 ΔP*_β-gt_*::*α-gt* and T4 ΔP*_α-gt_*::*β-gt*/P*_β-gt_*::*α-gt* in comparison to wild-type T4, upon infection in *E. coli* MG1655. The local virulence for each MOI was calculated from the corresponding growth curve in Figure 5C.

**Figure S13:**
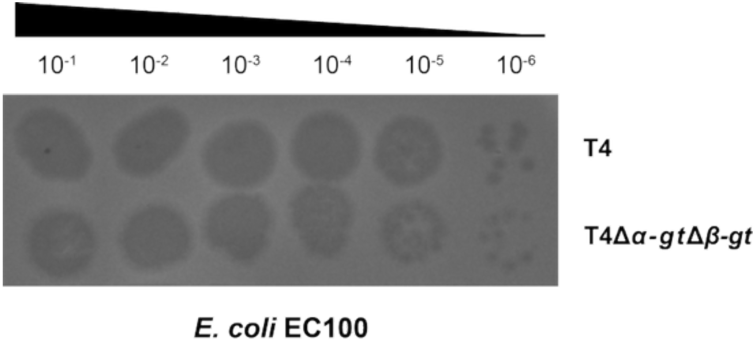
*E. coli* EC100 strain lacks type IV RM systems that target 5-hmC. Spot assay of serially diluted wild-type T4 and T4 1′*α-gt*1′*β-gt* on a lawn of *E. coli* EC100.

**Figure S14:**
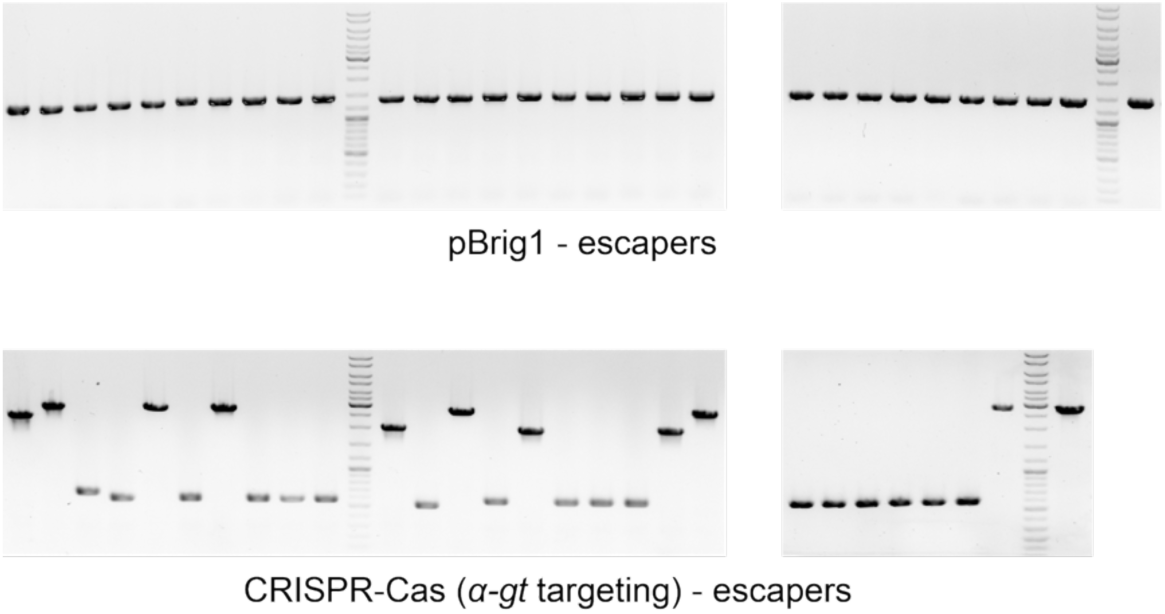
PCR analysis of the *α-gt* locus in phage T4 escapers generated post targeting by Brig1 (upper panel) or the CRISPR-Cas13b system (with a spacer complementary to the transcript of *α-gt*).

**Figure S15:**
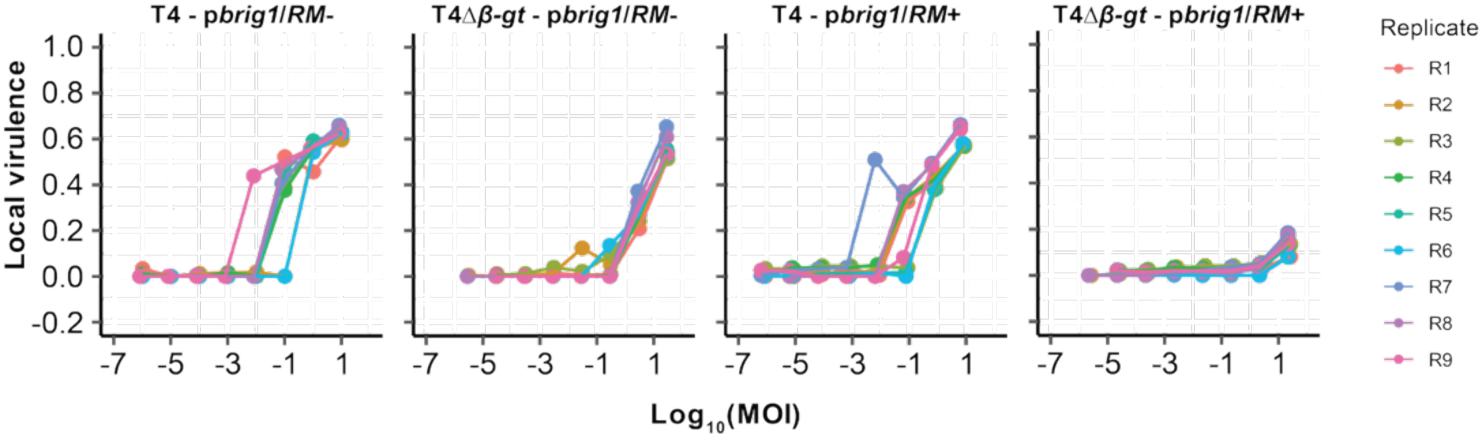
Local virulence corresponding to phage T4 and T41′*β-gt* upon infection in *E. coli* EC100 (RM-) and *E. coli* MG1655 (RM+). The local virulence for each MOI was calculated from the corresponding growth curve in Figure 6C.

## SUPPLEMENTAL TABLE LEGENDS

**Table S1**. Identification and analysis of α-GT, β-GT and β-glucosyl-HMC-α-GT homologs in phage genomes from the RefSeq and non-redundant (nr) NCBI GenBank database.

**Table S2**. Proteomic analysis of histidine-tagged ⍺-GT and β-GT pulldown samples.

**Table S3**. List of mutations in the *⍺-gt* locus from T4 escapers of Brig1 immunity.

**Table S4**. List of bacterial and phage strains, plasmids, and oligonucleotides used in this study.

